# Cortical thickness changes precede high levels of amyloid by at least seven years

**DOI:** 10.1101/2025.08.14.670398

**Authors:** James M. Roe, William J. Jagust, Susan M. Landau, Theresa M. Harrison, Håkon Grydeland, Maksim Slivka, José-Luis Alatorre-Warren, Pablo F. Garrido, Øystein Sørensen, Edvard O. S. Grødem, Tyler J. Ward, Esten H. Leonardsen, Tormod Fladby, Atle Bjørnerud, Kristine B. Walhovd, Anders M. Fjell, Didac Vidal-Piñeiro, Yunpeng Wang

## Abstract

Alzheimer’s disease (AD) is now defined based on its underlying brain pathology^1^, with the presence of amyloid (Aβ) plaques at high enough levels sufficient to warrant a diagnosis in the absence of cognitive symptoms. High levels of PET-detectable Aβ are widely thought to be the first imaging marker, with structural brain changes detectable on MRI scans thought to occur later. We combined 4570 longitudinal MRIs and 1684 Aβ PET scans from three cognitively healthy cohorts to test the difference in cortical thickness and its change between those that subsequently converted to be Aβ-positive or stayed Aβ-negative, using MRIs acquired exclusively in the years before conversion. We found those that subsequently developed elevated Aβ levels show both thicker cortex and less cortical thinning, even when the last MRI used to estimate their thickness trajectories was acquired at least seven years before conversion. Many effects remained when accounting for quantitative Aβ levels, suggesting some cortical thickness effects may be partly independent of Aβ. Differences in cortical thickness and its change between converters and Aβ-negative individuals showed moderate alignment with patterns of Aβ deposition, and the timing of thickness changes tracked the temporal progression of Aβ accumulation. Thus, if amyloid is AD^1^, we show that high levels of PET-detectable amyloid are not the first imaging marker of AD, as cortical thickness changes can be traced years before pathological amyloid. This has implications for understanding the sequence of events leading up to the earliest stages of AD.

## Introduction

As of July 2024^1^, the proposed consensus in Alzheimer’s disease (AD) research is that the presence of amyloid-beta (Aβ) plaques at high enough levels is sufficient to establish a diagnosis of AD^1^. Changes in Aβ as detectable by Positron Emission Tomography (PET) imaging are thought to be the first pathological marker of the disease and according to this view the biological manifestation of the disease itself^1–4^. Brain structural alterations detectable with MRI are thought to occur later. Indeed, studies on the temporal sequence of biomarker changes in the years before an AD diagnosis^5^ or expected onset of clinical symptoms^6,7^ suggest brain structural changes occur later than high levels of Aβ (i.e., Aβ-positivity), consistent with disease progression models of biomarker sequencing^8–11^. However, as the most common MRI-based marker of neurodegeneration, studies typically focus on measures of hippocampus^5–7^ or global brain volumes and their temporality within the disease process^8–11^. Whether wider brain structural changes may be evident in the years preceding high Aβ levels remains unknown, and few studies examine brain changes using MRIs acquired exclusively before Aβ-positivity. Consequently, it is unclear whether people who later go on to develop high Aβ show differences in brain structural markers in the years prior to becoming Aβ-positive.

Studies on the earliest stages of AD often compare brain measures between those with low Aβ levels and those with high Aβ absent other, later-occurring tau biomarkers^12–16^. Findings from these consistently suggest that distinct cortical changes may occur early relative to later in the disease process^12–17^. For example, a cross-sectional study found that global Aβ levels were positively correlated with cortical thickness in cognitively healthy adults with low tau, but negatively correlated with thickness in those with high tau^12^. Several other studies suggest that the relationship between Aβ and cortical measures may follow this biphasic model^13–16^, with a relatively thicker cortex in Aβ^+^ groups absent tau, followed by the expected more atrophy in those with both high Aβ and tau^12,14,15,17^. Specifically, studies suggest preclinical AD groups (Aβ^+^/T^−^) may show larger cross-sectional cortical measures^12,14,15,17^ and less longitudinal atrophy^14^ compared to Aβ^−^/T^−^ groups. Though reported regions differ, suggestions of a link between amyloid and cortical thickness have been found in at least five human datasets^12–14,17,18^, in mice engineered to overexpress the amyloid precursor protein (APP) gene^19^, and are hinted at in anti-Aβ drug trials — where the removal of Aβ seems to causally lead to more cortical thinning^20^.

Here, we combined longitudinal MRI data from three cognitively healthy cohorts (4570 scans from 1051 individuals, each with 2–14 timepoints) and focused on MRI scans collected in the years before individuals first showed Aβ-positivity. A total of 1,684 Aβ PET scans (691 individuals, 1–7 time points each) were used to define two groups: those who converted from Aβ^−^ to Aβ^+^, and those who remained Aβ^−^ across all scans. Quantitative measures of Aβ were also derived and included as covariates in subsequent analyses.

To examine brain structure before Aβ accumulation, we truncated each person’s MRI series so that the last scan included occurred at least X years before their first Aβ^+^ scan (X = 1-10 years). Using all available longitudinal MRI data, we then modelled cortical thickness trajectories across age with nonlinear mixed models. From these, we derived cortex-wide estimates of thickness and its change based solely on MRI data acquired ≥X years before the first Aβ^+^ scan observation. We compared these estimates between converters and Aβ^−^ non-converters, both with and without adjusting for quantitative Aβ levels. Finally, we replicated the analyses in a complementary sample using MRI data acquired exclusively in the years before a person was predicted to develop high Aβ levels.

We found that the cortex of those who subsequently become Aβ^+^ is both thicker and exhibits less longitudinal thinning compared to those with low Aβ, and the effects were detectable in MRI data acquired at least seven years before they converted to be Aβ^+^. Many of the observed cortical thickness effects persisted when accounting for quantitative Aβ levels, and thickness effects showed both moderate overlap with patterns of Aβ deposition and exhibited timing that tracked the temporal progression of Aβ. Thus, MRI markers of cortical thickness apparently precede pathological levels of Aβ accumulation on PET scans.

## Results

We gathered longitudinal MRIs from adults aged 30+ in samples that also had longitudinal Aβ PET scans, concatenating three densely sampled adult lifespan and ageing cohorts (LCBC, BACS, and ADNI [controls]). From these MRI data, we estimated individual-specific intercepts (baseline cortical thickness) and slopes (rates of change) using Generalized Additive Mixed Models (GAMMs) of age, adjusting for sex, scanner field strength, cohort, and intracranial volume (ICV; Methods). LCBC is comprised primarily of cognitively high-performing adults in the Oslo area of Norway, BACS participants are drawn largely from the Berkeley area of California, and from ADNI we included only individuals who were cognitively healthy across all diagnosis timepoints. For ADNI, we used only the MRI observations from the scanner field strength with the most timepoints per-person (or 3T when equal) to mitigate bias from scanner changes over time (Methods). This resulted in a total combined MRI sample of 4570 MRI scans from 1051 individuals (SI Table 1).

Of these, 691 had Aβ PET data, totalling 1684 scans. We used these to define two groups: those who converted from Aβ^−^ to Aβ^+^ and those who were Aβ^−^ on every available PET scan. For each converter we ensured the first PET timepoint was Aβ^−^ and the last was Aβ^+^. The time trajectories of the MRI and Aβ PET data described below are shown in Fig. 1A for converters (Aβ levels in SI Fig. 1; see SI Fig 2 for the Aβ^−^ group).

**Figure 1.**
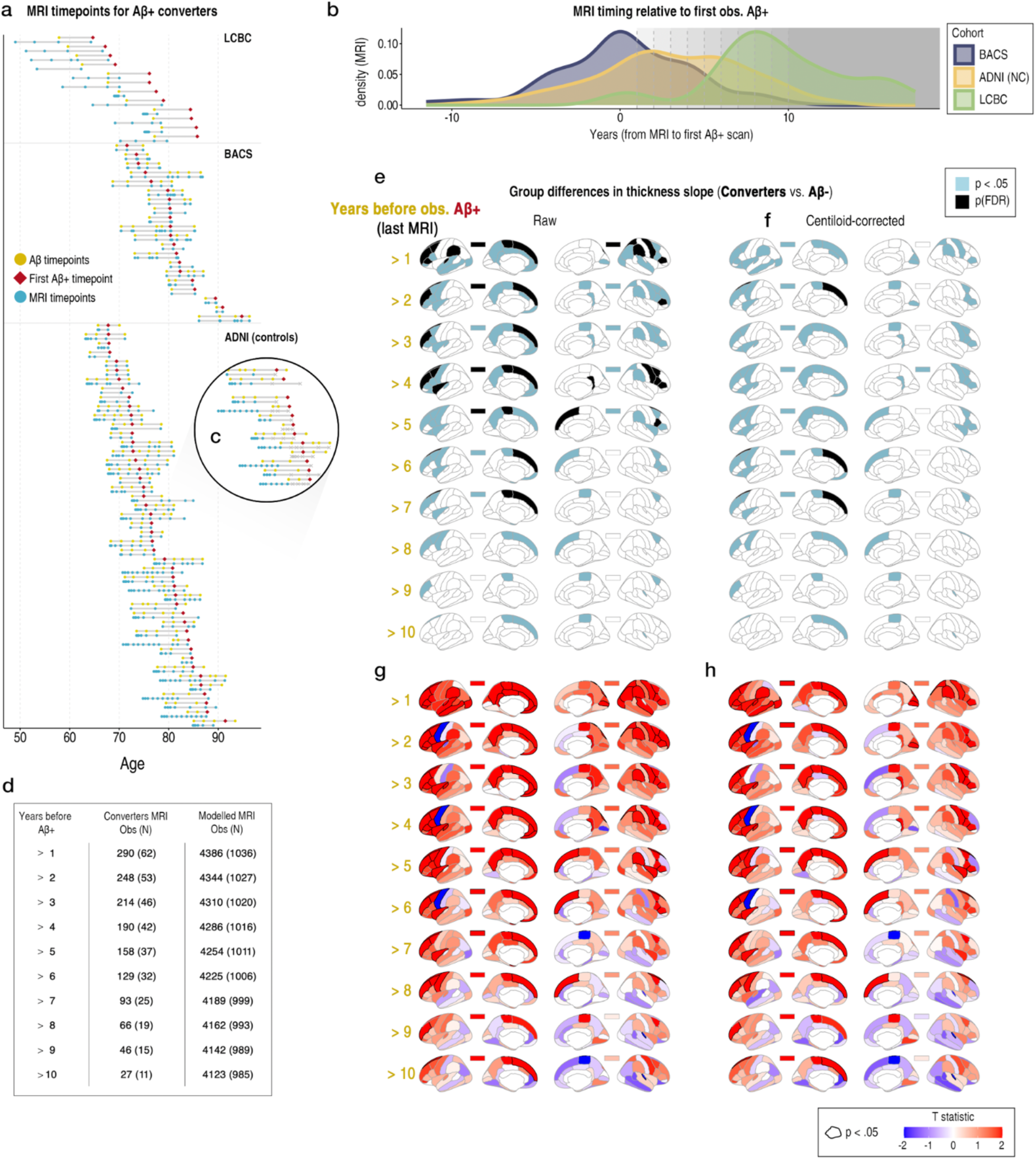
Analysis approach and differences in estimated slope of cortical thickness between converter and Aβ^−^ groups. **a** Time trajectories of Aβ PET/MRI scans for the initial 77 converters. The Aβ PET scans for each participant are in yellow and the first Aβ^+^ scan is depicted with a red diamond (later scans are also Aβ^+^). Below each is the MRI scans on that participant in blue. **b** The timing of MRI scans relative to observed Aβ-positivity shown as a density plot of the number of years from each MRI to the first Aβ^+^ scan. Colours depict cohort. Grey dashed lines represent the 1–10 year cutoffs applied, and shaded areas to the right of each line indicate the number of MRI scans included at each cutoff, ensuring the last MRI modelled was acquired at least X years prior to the first Aβ^+^ scan. Note that densities are approximate due to an additional requirement of ≥2 MRI time points at each cutoff (Methods). **c** The black circle highlights an example of the data truncation at a 4-year cutoff (MRI scans retained for analysis are in blue, while discarded scans—those within 4 years of the first Aβ^+^ scan—are marked with grey crosses). **d** The number of MRI observations and N in the mixed-model estimation at each time cutoff, shown for Aβ^+^ converters and the total sample modelled in the GAMMs. For this analysis, the number of scans and individuals (N) in the Aβ^−^ group remained stable at 1844 (411). See SI Fig. 6 for results when matching the Aβ^−^ group MRI trajectories on age and follow-up time. **e-f** Significant differences in the estimated slope of cortical thickness between Aβ^+^ converters and Aβ^−^ groups. Black parcels depict FDR-significant hits, whereas blue depicts significance at p<.05 (uncorrected). The rectangles above each brain hemisphere depict the results for mean cortical thickness. **e** “Raw” differences uncorrected for Aβ levels. **f** Differences after correcting for quantitative Aβ (mean centiloids across PET scans prior to the first Aβ^+^ scan). **g-h** Corresponding T maps (unthresholded).

**Figure 2.**
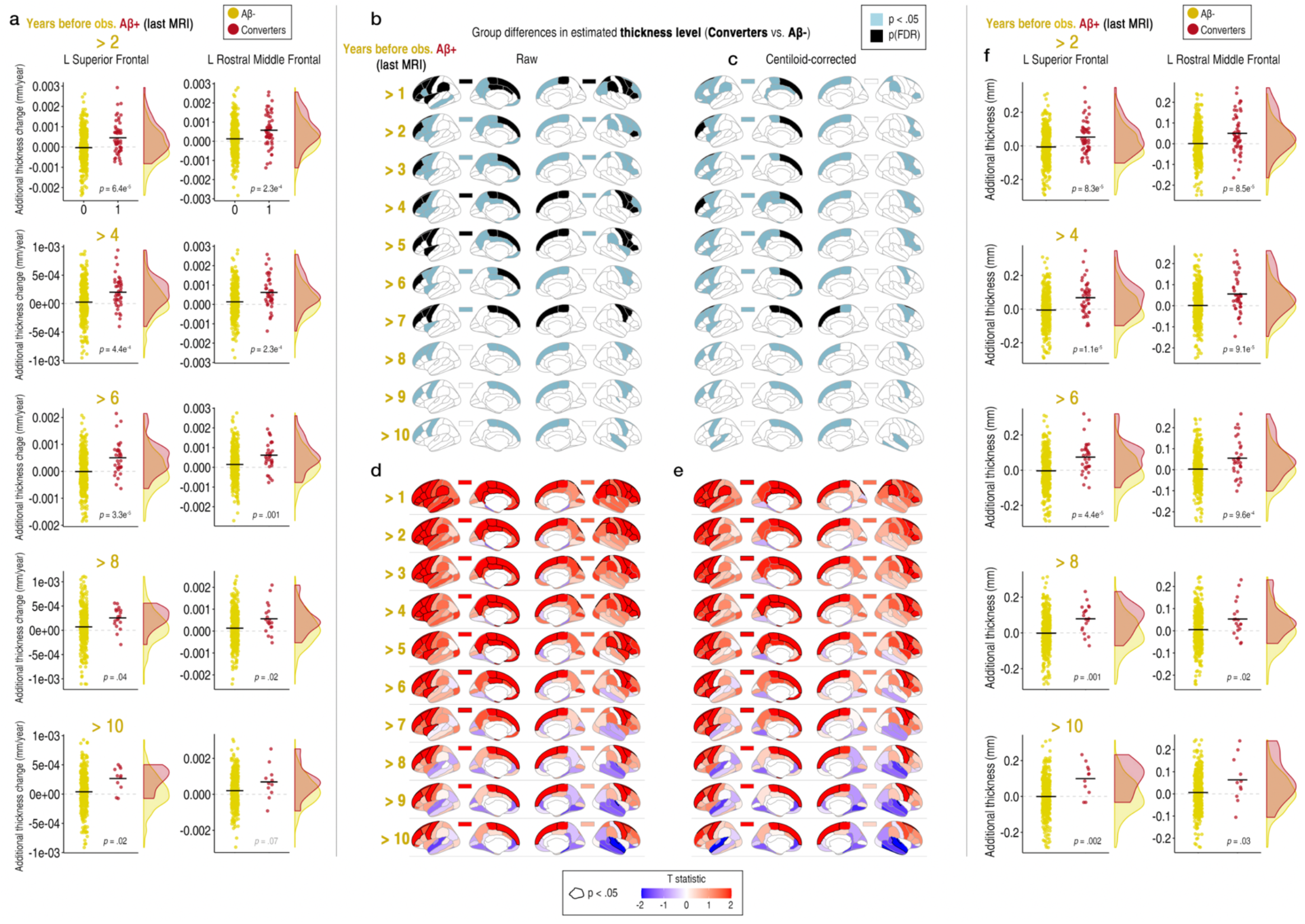
Differences in estimated cortical thickness level between converter and Aβ^−^ groups. **a** Differences for the models in Fig. 1E plotted for two example FDR-significant ROIs at four time cutoffs (estimated slopes). Datapoints corrected for sex, cohort, mean age, N timepoints, and interval between first and last MRI timepoint. The grey dashed line at 0 represents the average rate of change for age, which was estimated to be negative (SI Fig. 3), such that higher values indicate less cortical thinning in converters. **b-c** Significant differences in estimated cortical thickness level between Aβ^+^ converters and Aβ^−^ groups (intercepts; intercept-only models). Black parcels depict FDR-significant hits, whereas blue depicts significance at p<.05 (uncorrected). The rectangles above each brain hemisphere depict the results for mean cortical thickness. **b** “Raw” differences uncorrected for Aβ levels. **c** Differences after correcting for quantitative Aβ (mean centiloids across PET scans prior to the first Aβ^+^ scan). **d-e** Corresponding T maps (unthresholded). **f** Differences plotted for two example FDR-significant ROIs at four time cutoffs. The grey dashed line at 0 represents the average thickness level for age.

The converter group initially consisted of 77 individuals with 283 PET scans (2-7 timepoints; Fig. 1a) and a maximum of 487 MRI scans (2-14 timepoints; Fig. 1a). The Aβ^−^ group initially consisted of 412 individuals with 954 Aβ PET scans (1-7 timepoints; Aβ levels in SI Fig 2) and in total 1850 MRI scans (2-12 timepoints; one strong outlier was detected and removed from the MRI data; Methods). Note that we also included those with only a single PET timepoint in this group to maximise power. Individuals who started out Aβ^+^ were excluded from the Aβ PET analysis, but their MRI data were included in the total MRI sample.

For the main analysis, we calculated the time from each MRI to the first Aβ^+^ scan for each converter (Fig. 1b), then truncated the longitudinal MRI data by excluding scans collected within X years of their first Aβ^+^ scan (Fig. 1c). For each time cutoff (X = 1-10 years), MRI scans within that window were excluded, and the GAMMs re-run to yield change estimates at different distances from the observed conversion timepoint (see Fig. 1e for the number of MRIs modelled at each cutoff).

We then compared the model-estimated cortical thinning rates between converters and Aβ-participants. For each cutoff, we used the subject-specific slopes from the GAMMs as the outcome in linear models testing group difference in thickness change, adjusting for sex, cohort, a person’s mean age, number of MRI timepoints, and the time interval between the first and last MRI. The slopes reflect how quickly cortical thickness changed with age relative to the group’s average trajectory. A second set of models additionally adjusted for quantitative Aβ levels, using the mean centiloid value across an individual’s Aβ PET timepoints. For converters, only PET scans prior to the first Aβ^+^ scan were included in the mean.

Fig. 1d-e shows the group-difference results for the model-estimated slope in cortical thickness at each time cutoff. Black parcels indicate a significant difference after correction for the False Discovery Rate (FDR; 70 tests per map), whereas blue parcels denote significance at p<.05 (uncorrected). Because statistical power varied across time cutoffs (Fig. 1f), parcels that were FDR-significant at any cutoff were considered significant at less-powered cutoffs at p<.05 (uncorrected). We found all FDR-significant differences showed positive effects, and the results across maps at varying time cutoffs were highly consistent (Fig. 1e). Because the average rate of change across age was estimated to be negative (SI Fig. 3; i.e., cortical thinning is expected), the positive differences indicate less cortical thinning in converters. Particularly in frontal cortex we observed many significant regions where rates of thinning were lower in converters. Importantly, these were in regions that remained significant also at later time cutoffs, i.e. further away from the first positive PET, despite the lower sample sizes (Fig. 1d); in left superior frontal cortex, the data indicated a significant difference (p<.05 [uncorrected]) in estimated thickness change was detectable even when the last MRI in the converter group was taken 10 years prior to their first Aβ^+^ scan observation. Although some effects were attenuated, many remained also after adjusting for differences in Aβ levels (mean centiloids; Fig. 1f), suggesting they were not solely driven by emerging Aβ levels. A resampling-based analysis confirmed the reported significance results were robust to sample variations and not driven by outliers (SI Fig. 4), and permutation-based analyses confirmed the results were unaffected by group-differences in variance of the estimates (SI Fig. 5).

The positive group differences in estimated thickness slope suggest that individuals who later go on to become Aβ^+^ show less cortical thinning compared to Aβ^−^ individuals. That is, although both groups show overall negative change (SI Fig. 3), the results indicate Aβ^+^ converters show less negative change over time, i.e., less age-related thinning (see plots in Fig. 2a).

### Sensitivity analyses

To confirm that group differences were not driven by longer or denser MRI follow-up in the Aβ^−^ group, we repeated the analyses using age- and follow-up-matched Aβ^−^ samples (Methods), and the results remained robust, indicating that the observed differences in cortical thinning were not explained by imbalances in longitudinal data (SI Figs. 6-7)

To rule out the possibility the group differences were driven by the field strength of the MRI scanner, we reran the analysis retaining only MRI scans acquired at 1.5T, with and without matching groups on age and follow-up data. Again, the results were robust, suggesting the group differences are likely not an artefact caused by differences in scanner field strength (SI Fig. 8; SI Fig. 10).

To address potential confounding between slope and intercept in the random-effects models, we reran the analyses using only random intercepts (i.e., thickness level), representing baseline thickness relative to age without estimating change. The same regional group differences emerged, indicating that individuals who later became Aβ^+^ already showed a thicker cortex—particularly in frontal regions—up to 10 years before their first Aβ^+^ scan (Fig. 2; SI Fig 9; SI Fig. 11), and that the contributions of intercept and change to the estimated slopes were not fully disentangled in our initial model.

To help disentangle the contributions of level and change, we ran alternative GAMMs to better statistically separate the slope and intercept in the random effects, albeit with reduced statistical power (tensor smooth interaction models; Methods; SI Fig. 12). To further rule out potential confounds when modelling change, we did this using only 1.5T scans and matching the groups on age and follow-up data, and both with (SI Fig. 14) and without adjusting for thickness level (random intercepts; SI Fig. 13). The results again remained robust, with significantly reduced longitudinal thinning in converters evident in MRI data acquired more than 8 years before the first Aβ^+^ scan—confirming the results from our initial model indeed reflect both cortical change and level effects. When additionally adjusting for centiloid values, although some regions remained significant, the attenuation of effects was more pronounced, suggesting the change effect was less statistically independent of emerging Aβ. More manual calculations of linear change also confirmed the longitudinal effect as identified and amplified via the mixed-model approach (SI Fig. 15).

Because the MRI data was processed longitudinally, each timepoint’s processing incorporated information from all of a person’s timepoints. To ensure this did not introduce bias—where earlier brain measures could be influenced by later ones—we also reran the initial main analysis using cross-sectionally processed brain measures, finding near-identical results (SI Fig. 16).

### Additional analyses

To test overlap between the observed thickness effects and Aβ deposition patterns, we ran spatial correlations using permutation-based spin tests^21^ after computing the following three maps: 1) the group-difference in standard uptake value ratio (SUVR) between converters and the Aβ^−^ group (Methods), 2) the average SUVR map from all Aβ^+^ scans in an independent ADNI sample with an MCI or AD diagnosis (1314 PET scans; N = 734; Fig. 3a), 3) the group-difference in SUVR between diagnosed Aβ^+^ individuals and the Aβ^−^ group. Note that although the mean uptake in superior frontal regions appears low in both converter and diagnosed Aβ^+^ groups (Fig. 3a,b), the difference in its uptake with the Aβ^−^ group was nevertheless large (Fig. 3c,d). In a multiverse-style approach^22^, we ran spatial correlations with the effect maps derived across six analyses (main and sensitivity; Fig. 3e,f).

**Figure 3.**
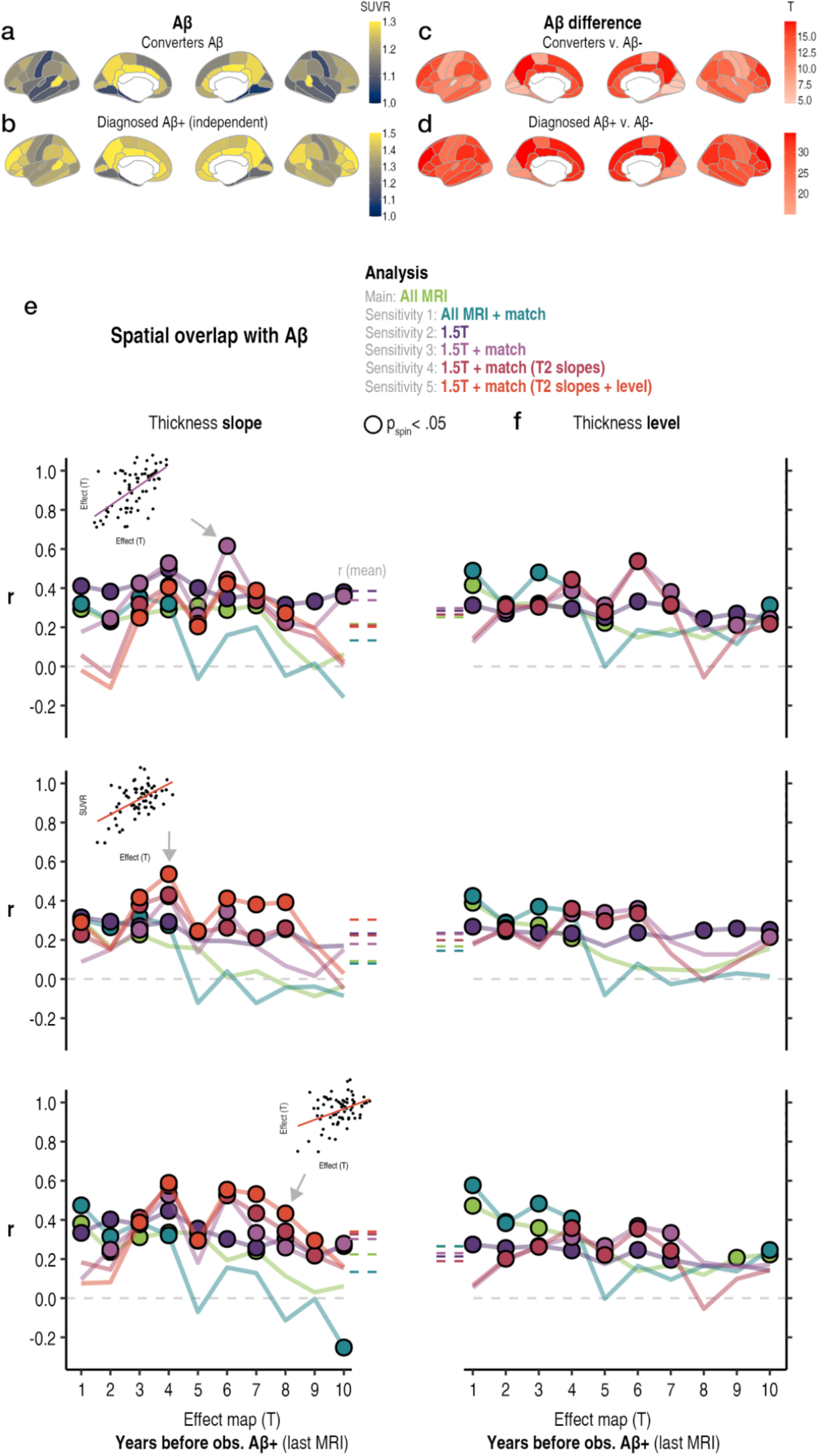
Spatial overlap with Aβ. **a** Average SUVR map in the converter group (top) and **b** an independent sample of Aβ^+^ individuals with an MCI or AD diagnosis (Methods). **c** Effect size for the group-difference in SUVR (T) between converters and the Aβ^−^ group (top), and **d** for the group-difference between diagnosed Aβ^+^ individuals and the Aβ^−^ group (bottom; both tracer-adjusted, all p[FDR]<.05). **e-f** Spatial correlations between the cortical thickness effect maps at each time cutoff for six analyses, and the maps in **c** (top), **b** (middle), and **d** (bottom). Circles depict significant permutation-based spatial correlations (p_spin_<.05). **e** Correlations with the estimated slopes. **f** Correlations with the estimated intercepts. The mean correlation across all time cutoffs per analysis is shown as a dotted line to the left/right of each plot.

There was clear overlap between the observed thickness effect maps and spatial patterns of Aβ deposition, with 110 out of 180 (70.1%) tests exhibiting significant overlap with the estimated slopes (p_spin_ < .05; Fig. 3), and 91 of 150 significant tests for estimated thickness level (intercepts). Although the global overlap was not especially strong (Fig. 3), it was generally stronger for analyses that only incorporated 1.5T scans, supporting our analytical choices. Moreover, although models that tested only for differences in thickness level showed some spatial overlap (Fig. 3e), models that isolated change generally showed strongest overlap (in two of three cases; Fig. 3e) and yielded the highest number of significant correlations with Aβ deposition patterns.

Together, these results support that 1) the reported thickness effects show moderate overlap with Aβ deposition patterns, 2) our change models captured true differences in thickness change, and 3) that a mixture of preexisting and change effects may underlie the observed group differences in cortical thickness.

While the current time cutoffs allow for the most statistically powered tests, they are based on when Aβ positivity was first observed—not when it was actually reached. To address this, we estimated each converter’s predicted age at Aβ positivity based on their centiloid trajectories (Methods; Fig 4a).

**Figure 4.**
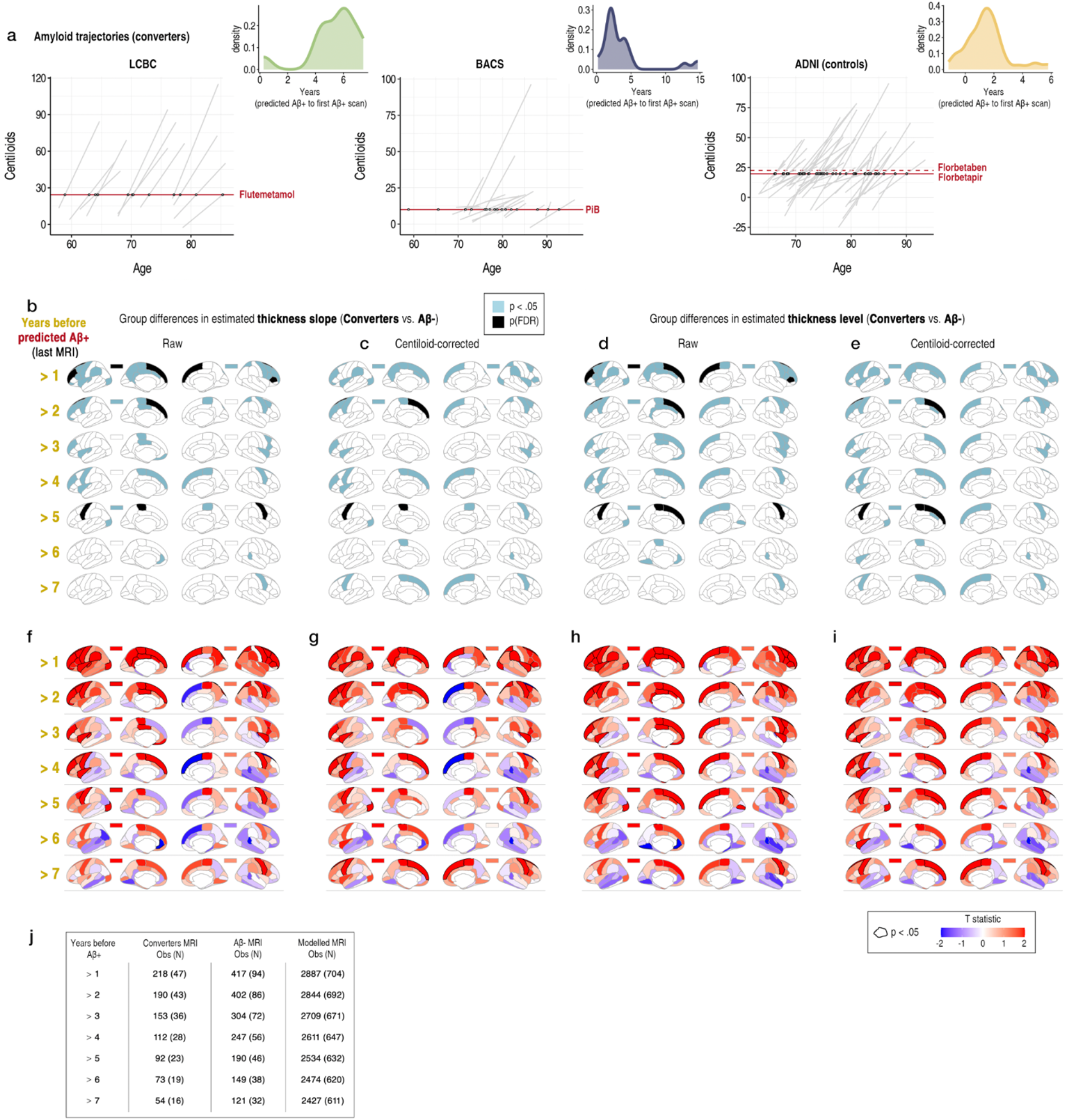
Differences in estimated cortical thickness between matched converter and Aβ^−^ groups using predicted age at amyloid positivity. **a-c** Amyloid trajectories in centiloids for the Aβ^+^ converters in each cohort (linear model across PET timepoints; raw data in SI Fig. 1; see SI Fig. 19a for the first Aβ^+^ scans in each cohort). The sample-specific threshold for Aβ positivity is shown as a solid red line (Methods), and predicted age on crossing the threshold is shown by a white dot (note we used the Florbetapir threshold in ADNI which was lower). We used this to ensure the last MRI included in the analysis was acquired 1-7 years before the participant was predicted to become Aβ^+^ (Methods). Inset plots show the time from predicted to observed Aβ positivity (i.e., the first Aβ^+^ scan) for each cohort. Note that as LCBC had only two PET timepoints, more individuals were predicted to cross the Aβ^+^ threshold several years before they were observed Aβ^+^. Note also that two individuals in BACS were predicted to cross the threshold at very low ages, likely attributable to measurement error (SI Fig. 24). **b-c** Significant differences in cortical thickness slope, with and without adjusting for amyloid levels (mean centiloids across PET scans prior to the first Aβ^+^ scan; see also SI Fig. 19). **d-e** Significant differences in cortical thickness level (intercept-only models), with and without adjusting for amyloid. Black parcels depict FDR-significant hits, whereas blue depicts significance at p<.05 (uncorrected). The rectangles above each brain hemisphere depict the results for mean cortical thickness. **f-i** Corresponding T maps (unthresholded). **j** The number of MRI observations and N in the mixed-model estimation at each time cutoff, shown for both Aβ groups and the total sample submitted to the GAMMs. Groups were matched on age and follow-up data (Methods).

Note that, because LCBC had only two PET timepoints, many individuals were predicted to have crossed the Aβ^+^ threshold several years before they were observed Aβ^+^. To ensure that LCBC converters who may have been Aβ-positive for longer did not disproportionally affect the results, we first excluded their data and reran the analysis. Group differences in thickness remained highly similar in BACS and ADNI converters (where the first Aβ^+^ scan typically occurred soon after predicted Aβ-positivity; SI Fig. 17a), suggesting the results were not driven by LCBC converters who had likely been Aβ-positive for longer (SI Fig. 17). This was also the case when only using 1.5T scans (SI Fig. 18). Notably, similar differences in frontal cortex were detectable in BACS and ADNI MRI data acquired up to 9 years before the first Aβ^+^ scan (SI Fig. 17).

We then re-ran the analysis using time-to-predicted Aβ positivity as the truncation metric (1–7 years before predicted Aβ^+^ onset). To boost power to detect the effects in this further reduced sample (see Fig 3j; at the 7 year cutoff, N converters = 16)—and having ruled out scanner field strength as a confound—we ran the GAMMs using both 1.5T and 3T scans, and using our initial model (see also SI Fig. 20 for tensor smooth interaction models using 1.5T scans). The earliest MRI from converters was acquired 11.8 years before conversion. The results confirmed that differences in cortical thickness (slope and level) between Aβ^+^ converters and Aβ^−^ individuals were detectable also when the last MRI was acquired more than 7 years before a person became Aβ^+^ (Fig. 4).

As a final step, we tested whether there is more thickness change in regions with higher amyloid accumulation, and the temporal correspondence between thickness changes and Aβ changes in regions with significant thickness effects (Methods). Here, we used all MRI scans in converters (i.e., also those acquired after conversion), and ran GAMMs of time-to-predicted Aβ positivity on thickness in this group only (Methods).

The thickness trajectory in high Aβ regions was significantly different than low Aβ regions (p = 5.7^-5^), and the most profound thickness increases occurred around and prior to conversion (Fig. 5a-b). In other words, Aβ^+^ converters showed a relative intraindividual thickening in high Aβ uptake regions, closely associated with Aβ conversion time.

**Figure 5.**
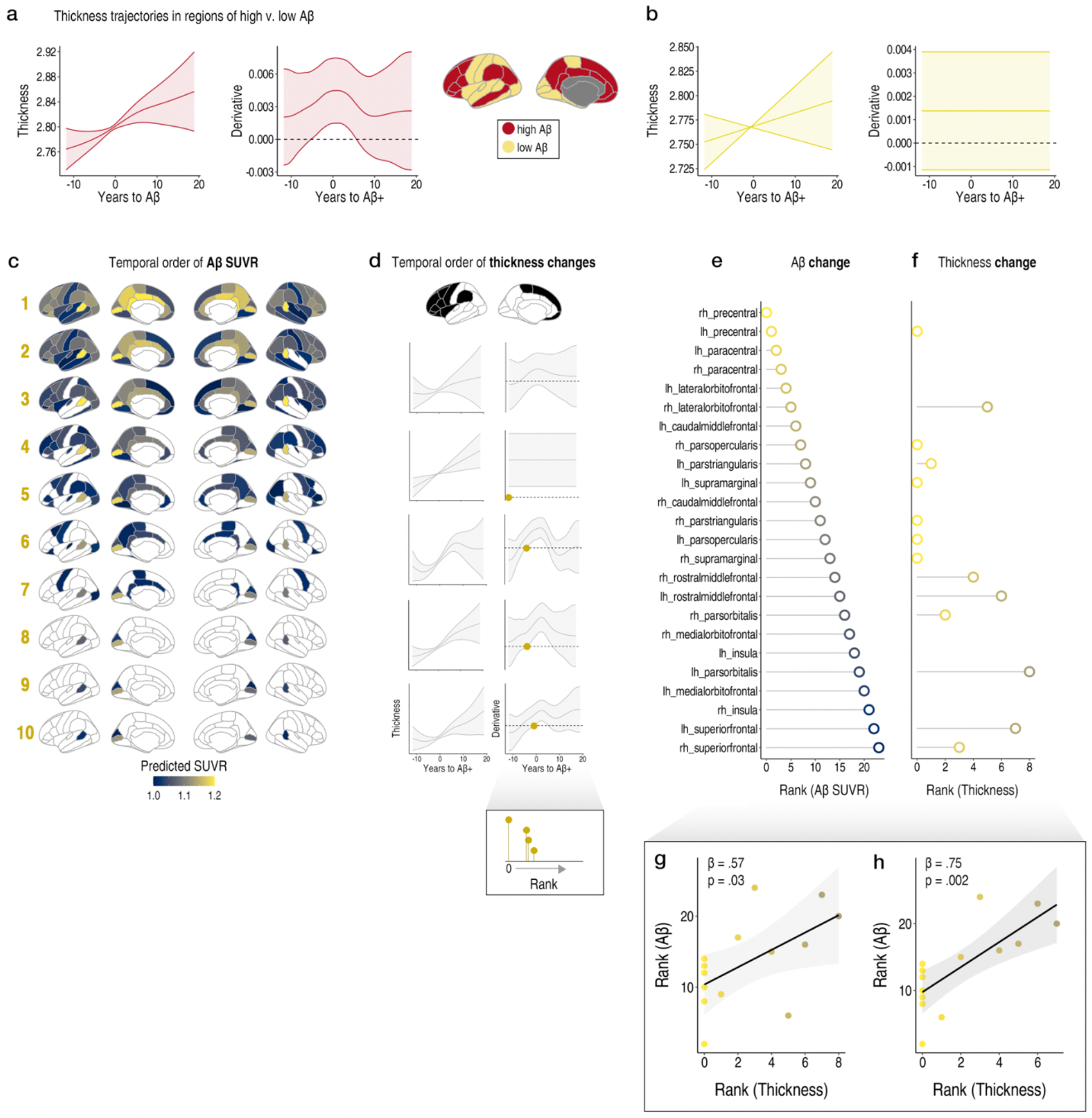
Timing of Aβ-positivity and cortical thickness changes. **a** Average thickness trajectory and rate-of-change (i.e., derivative) in regions of high and **b** low Aβ deposition as a function of years to predicted Aβ-positivity. Bands represent 95% confidence intervals (CIs). **c** Predicted Aβ SUVR as a function of years to predicted Aβ-positivity. The cortical maps are thresholded at 1.0, indicating uptake greater than the reference region (cerebellar grey). The maps give an indication of the temporal order of Aβ accumulation and were used to derive the rank order of Aβ SUVR changes (Methods). **d** Estimating the temporal order of thickness changes within regions showing significant differences in thickness between converter and Aβ^−^ groups (Methods). Example plots of the regional thickness trajectory and rate-of-change (i.e., derivative) for five regions. The point at which the derivative CI excluded zero was used to estimate the rank order of thickness changes (marked with gold dots; dashed lines at zero). **e** The rank order of Aβ SUVR changes and **f** thickness changes. Missing data points reflect regions where the derivative CI did not exclude zero and thus could not be ranked. **g** The correlation between the temporal order of Aβ change and thickness change (derived using CIs). **h** The correlation between the temporal order of Aβ change and thickness change (derived using standard errors).

Finally, we estimated the spatiotemporal trajectory of Aβ (linear mixed models of time-to-predicted Aβ positivity on Aβ SUVR; Methods). The maps in Fig. 5c illustrate the Aβ trajectory based on uptake exceeding the reference region (note that regions with low mean uptake but large relative differences in Aβ^+^ groups will be underrepresented; e.g., superior frontal regions; Fig. 3a-d). In regions showing significant group-differences in thickness, we derived the temporal order of Aβ changes (Fig. 5e; based on the point where predicted SUVRs exceeded reference region uptake) and thickness changes (Fig. 5d; based on the point where the CI of the derivative excluded zero). The correlation between the rank order of Aβ changes and thickness changes was significant (r = .57, p = .03; Fig. 5g), and stronger when applying a more relaxed threshold to define thickness changes (standard errors; r = .75, p = .002; Fig. 5h). Notably, many regions estimated to show the earliest Aβ changes showed linear rates-of-change of time-to-Aβ on thickness (i.e., the derivative CI excluded zero from the outset and thus were ranked as zero; Fig. 5e-f), suggesting rising thickness levels many years prior to Aβ-positivity. Overall, these results suggest the timing of thickness changes largely parallels the temporal progression of Aβ accumulation, and that regions showing the earliest thickness differences between converters and Aβ^−^ groups tend to be those where amyloid first accumulates.

## Discussion

Our findings suggest that structural brain changes precede pathological levels of amyloid by several years, with implications for how we should understand the sequence of events leading to Aβ-positivity. Using MRI data acquired in the years before high Aβ levels, we found those who subsequently became Aβ-positive have an apparently thicker cortex and show less longitudinal thinning, compared to those with low Aβ levels. Differences in cortical thickness were detectable in MRI data acquired at least seven years prior to individuals reaching Aβ levels consistent with a biomarker-defined AD diagnosis^1^. By showing that cortical thickness changes precede high levels of Aβ, this work adds to our understanding of the brain changes that unfold before detectable amyloid pathology.

The results were robust to analysis variations to rule out confounding related to MRI scanner, differences in longitudinal follow-up, and MRI processing. In our main models, we observed strong evidence that subject-level estimations of both the change and level of cortical thickness differed significantly between individuals who later became Aβ^+^ compared with Aβ^−^ individuals, and throughout frontal cortex (Fig. 4; SI Fig. 19). Many of these remained after adjusting for Aβ levels, suggesting some of the differences may be at least partly independent of emerging amyloid. However, because individuals with greater cortical thickness for their age also tended to show slower thinning over time, the contributions of level and change could not be fully disentangled. While this correlation is expected, it means that the observed group differences likely reflect a combination of both a thicker cortex and reduced cortical thinning—i.e., relative cortical thickening on top of ageing effects—among those who later develop high Aβ.

Where we could directly attribute the effects to longitudinal change, we found less longitudinal thinning in Aβ^+^ converters, and in the same frontal cortical regions, supporting the interpretation that the differences reflect both greater thickness and reduced age-related thinning. Notably, less longitudinal thinning was evident both when the last MRI was acquired at least 8 years before the first Aβ^+^ scan (Fig. 3), and several years prior to conversion to Aβ-positivity (SI Fig. 20). Note the former does not necessarily exclude Aβ^+^ MRI timepoints, whereas the latter does (Limitations). This suggests the effect can be detected in MRI data stretching back further in time (the earliest MRI was acquired 11.8 years before conversion). Interestingly, adjusting for Aβ levels most strongly attenuated the group differences in models that isolated the change effect. This suggests the longitudinal effect—less thinning in Aβ^+^ converters—may to a greater extent reflect and be captured by emerging Aβ pathology. However, it is also possible differences in cortical thickness level may reflect earlier change. Cortical thickness effects showed moderate but clear overlap with underlying Aβ deposition patterns, and we observed slightly higher overlap for the maps reflecting the longitudinal group-difference in thinning (Fig. 3). Critically, we found evidence of more thickness increase in high Aβ regions, and the timing of relative thickness increases paralleled the spatiotemporal progression of Aβ. In particular, several regions exhibiting earlier thickening also showed signs of earlier Aβ deposition (Fig. 5). Together, these results implicate both less age-related thinning, and possibly preexisting differences reflecting a thicker frontal cortex, in those that later become Aβ^+^. This contrasts with a common view on the temporal order of imaging markers, according to which Aβ imaging markers precede changes detectable on MRIs^4,6–10,23^.

The results could be interpreted within a biphasic model^15,16,24,25^, whereby Aβ deposition may lead to an initial cortical thickening—which against a background of dominant ageing changes would manifest as reduced cortical thinning^26^—before the development of later-occurring tau pathology which drives faster thinning^27,28^. Indeed, studies on preclinical AD suggest a link between Aβ and larger cortical measures in the earliest disease stages, with cross-sectional studies suggesting Aβ^+^/T^−^ groups show thicker cortex compared to Aβ^−^/T^−^ groups^13–16,18,29^, and that Aβ levels correlate positively with thickness in those with high Aβ absent tau^12^. In line with our findings, Pegueroles et al. reported less longitudinal thinning in Aβ^+^/T^−^ groups over 2-years in medial frontal regions^14^, and Harrison et al.^12^ found positive Aβ-thickness correlations in lateral frontal regions where we find thicker cortex in the earlier MRIs of converters. Note while some of the data here overlaps with these studies, we also confirm similar effects in the independent LCBC cohort (see SI Fig. 21). The effects also broadly align with greater cortical volumes reported in Aβ^+^/T^−^ individuals in the ALFA cohort^18^. Others have reported thicker cortices in Aβ^+^/T^−^ in temporoparietal regions that seem less consistent with our results^15,16,25^ – which mainly implicate frontal cortical regions – though it is possible this may reflect a temporal ordering to the thickening phase as Aβ accumulation progresses that would not be detected by our analysis.

In these studies, the groups in question had already developed high Aβ levels (i.e., were Aβ^+^/T^−^). Thus, we extend this earlier work to show that a relative cortical thickening phase in Aβ^+^ converters is detectable in MRIs acquired many years before they develop pathological Aβ levels. Several possible explanations could account for this. First, a thicker cortex in the years leading up to high Aβ might reflect an immune response to accumulating, sub-threshold Aβ deposits^30^, which is a process thought to span over two decades^5,23^. This view that early cortical thickness changes reflect neuroinflammation has some, albeit indirect, support from studies reporting associations between CSF markers of microglial activation and cortical structure in cognitively normal adults^18,31^, as well as a small study linking a PET-based marker of astrocytosis to cortical thickness in autosomal dominant AD^24^. That neuroinflammatory processes may be triggered prior to Aβ plaque formation also has support from animal models reporting increased glial recruitment and upregulation of inflammatory markers before plaques emerge^32,33^. Second, the presence of space-occupying Aβ plaques alone could be enough to account for concomitant increases in cortical thickness measured on MRIs^12,20^. This aligns with the overlap between thickness effects and Aβ accumulation maps, but the moderate spatial correlation and persistence of effects after adjusting for Aβ suggests it is unlikely to be the main explanation for the observed thickness differences. Third, our data are also compatible with preexisting effects of higher cortical thickness in those that subsequently develop high Aβ levels, which may underlie some of the group differences. Indeed, when only accounting for differences in thickness level (i.e., intercept), we found we could recapitulate many of the observed regional differences (Fig. 2; Fig. 4). However, it remains unclear to what extent intercept effects may also reflect change, as subtle and difficult-to-estimate structural changes will also influence estimates of thickness level, which are more robust, easier to quantify, and less noisy. Still, it is possible a thicker frontal cortex in individuals who later become Aβ^+^ may arise during development, possibly consistent with a study reporting less developmental cortical thinning in young mutant mice engineered to overproduce Aβ^19^. Alternatively, a thicker frontal cortex in those who later become Aβ^+^—yet remain cognitively healthy—could reflect a preexisting protective factor, consistent with a “brain reserve” account in which greater brain structural resources help buffer against the consequences of brain pathology (see Limitations). A fourth possibility—that cortical thickness increases represent an earlier biological process to which Aβ accumulation is a response, should also not be ruled out.

The first two explanations have recently been invoked to explain the finding from anti-Aβ drug trials that removing Aβ leads to greater observed brain volume loss in treatment groups, especially in the cortex^20^. By showing that higher cortical thickness—in part reflecting a relative cortical thickening—is evident in the years leading up to high Aβ levels, our study may help shed light on this phenomenon in human trials and inform the interpretation of MRI-based endpoints used to assess drug efficacy.

An intriguing aspect of our findings is the absence of amyloid–thickness relationships in the medial or lateral temporal lobes—regions vulnerable to the most degeneration in both ageing and AD and strongly linked to cognitive decline in each^34^. Studies have reported a similar lack of association in cognitively normal older adults, suggesting relative preservation of these regions may contribute to preserved cognition despite accumulating pathology^35^. This may help explain the dissociation between Aβ accumulation and cognitive decline^34,36^, and aligns with the idea that the temporal lobe memory network is vulnerable both to normal ageing and neurodegenerative processes, though the underlying mechanisms may differ. It therefore remains uncertain to what extent the cortical differences we observe years before amyloid positivity reflect early risk factors for cognitive decline.

Our study has several strengths. We leveraged all available longitudinal MRIs and the power of mixed models to estimate subject-specific trajectories in the subset of individuals that formed our Aβ PET groups. This statistical approach to estimating subject-specific effects benefits from as many longitudinal observations as possible, is psychometrically superior to more manual calculations of change, and less sensitive to outliers due to the shrinkage effect^37,38^. Briefly, this reduces the influence of extreme data points by estimating random effects based on a probability distribution informed by the full dataset, which becomes more robust with more longitudinal data, leading to more reliable estimates. Another strength is that we defined our groups using all available Aβ PET scans from individuals with longitudinal MRIs who remained cognitively healthy. We also capitalized on the opportunity to examine longitudinal MRIs acquired in the years before a biomarker-defined event in longitudinal Aβ PET data. In doing so, our study yields unique insights into the temporal dynamics of imaging markers in preclinical AD.

There are also limitations. First, to maximize power for detecting effects, we first applied a cutoff for MRI inclusion based on the number of years before the first Aβ^+^ PET scan observation. However, these time cutoffs can be misleading, as some individuals seemingly converted to Aβ-positivity up to 7 years before their first Aβ^+^ scan (Fig. 4a). This was most evident in the LCBC dataset which included only two PET timepoints (see SI Fig. 17–18 for comparable results excluding LCBC). As such, the initial cutoffs do not necessarily imply the MRI sample was entirely free of Aβ^+^ individuals—though their proportion decreases at later time cutoffs. Alternative approaches using time-to-Aβ-positivity use subject-level predictions to estimate the point of Aβ positivity (as in Fig. 4A) then infer timing from group-level predictions across aligned data^27,39^. Our approach based on only analyzing MRI scans acquired at least X years before an indicator of Aβ-positivity is thus fundamentally different, and in our initial models we retained as much power as possible to test it. As can be seen, applying the more precise cutoff based on years-to-predicted Aβ-positivity substantially reduced power (Fig. 4j), but nevertheless allowed us to confirm the effects we found in more powered analyses (Fig. 4). Hence, to avoid ambiguity, we refer to years before observed/predicted Aβ-positivity accordingly. Second, we included those with only a single PET timepoint in the Aβ^−^ group, such that some individuals sorted into the Aβ^−^ group may convert to Aβ^+^ had additional PET scans been available. However, as our goal was to compare retrospective MRI data between groups defined by their subsequent PET data, we reasoned this limitation will almost always apply to the Aβ^−^ group, and accepted the tradeoff to maximize power by leveraging more of the PET/MRI scans. Still, if some converters were misclassified into the Aβ^−^ group, it is possible this may have only attenuated the observed group differences. Third, we did not include tau biomarkers. However, the cornerstone of the amyloid hypothesis is that tau occurs downstream from Aβ, which drives subsequent tau that is the proximal cause of brain and cognitive losses^4^. Indeed, an abundance of evidence shows tau strongly predicts brain loss and cognition^27,40– 42^, whereas Aβ is not strongly associated with either^40,41,43^. Furthermore, studies that stratify on both Aβ and tau confirm tau only modifies Aβ-structure associations in the expected negative direction^13–16,18,29^. Thus, if Aβ precedes tau and our MRI data precedes high Aβ, then tau is unlikely to be a strong driver of our reported effects—which were all positive. Fourth, an Aβ^−^ scan doesn’t imply the complete absence of Aβ plaques, only that the existing plaque burden may be below the current detection threshold^1^. Though we additionally considered quantitative Aβ levels according to current best practices^44^, the results could vary depending on the applied threshold. However, any such variation is likely to be minimal, as we demonstrate our findings are robust across many analytic specifications, drawing influence from multiverse-style analyses that emphasize the consistency of statistical results across analysis variations^22^. Fifth, we note that differences in apparent thickness may rather reflect biologically meaningful differences in intracortical myelination rather than thickness per se, as MRI-based delineations of the cortical ribbon will be influenced by many factors. Sixth, we included only cognitively healthy individuals in our main analysis. This choice was made to reduce confounding by the presence of other biomarkers or pathologies, but also means the Aβ^+^ converters represent individuals who became Aβ^+^ while remaining cognitively healthy. As a result, some of the observed effects could reflect a greater capacity to tolerate pathology in converters (i.e., brain reserve). That said, one could also argue the Aβ^−^ group is similarly defined by resilience—specifically, the ability to age without developing Aβ pathology. In this sense, both groups represent different forms of successful ageing—one in the presence of pathology, the other in its absence—and future research is needed to clarify whether or to what extent the effects reported here may reflect preexisting reserve factors. Relatedly, the cognitively healthy samples included here are unlikely to be representative of real-world clinical samples, because longitudinal studies inevitably recruit and culminate in unrepresentatively high-performing samples. Still, a key tenet of the amyloid hypothesis is that Aβ is specific to AD^1,45,46^. Hence, according to new biomarker-based definitions^1^, the converter group here developed AD during the course of the study, despite their cognitive health. In defining our groups, we adopted these new criteria to test whether we could find an earlier imaging marker of biological AD.

In conclusion, we show that differences in cortical thickness are detectable in those that later develop high Aβ levels in MRI data acquired at least seven years before they crossed the threshold for amyloid positivity. Specifically, individuals who later become Aβ^+^ show both thicker cortex and less longitudinal thinning evident in retrospective MRIs acquired prior to Aβ^+^ conversion, compared to Aβ^−^ individuals. Our results have implications for understanding the sequence of events leading up to Aβ-positivity, showing cortical thickness effects many years prior to previous assumptions

## Methods

### Participants

We combined the longitudinal structural brain MRI data from three densely sampled adult lifespan and ageing MRI cohorts (LCBC, BACS, and ADNI [controls]). We included only cognitively healthy, non-diagnosed adults, and each cohort had Aβ PET data for a subset of individuals. We gathered all available longitudinal MRIs, and all available Aβ PET scans from the cognitively healthy adults in each sample. To help ensure reliable MRI change estimates, participants with <0.5 years of follow-up data were excluded. The total combined MRI sample initially comprised 4570 MRI scans from 1051 individuals (2-14 timepoints [median = 4]; follow-up range = 0.5– 16.0 years, mean age [SD] = 72.1 [11.4]; 607 females). Of 691 individuals that also had Aβ PET scans, we used a total of 1684 Aβ PET scans to define two groups: those who converted to be Aβ^+^ from an earlier Aβ^−^ scan (converter group) and those who were deemed Aβ^−^ on every available PET scan (Aβ^−^ group). An initial 77 individuals were found to convert from an Aβ^−^ scan to an Aβ^+^ scan at a later point in time (across 283 Aβ PET scans; 2-7 timepoints; Fig. 1A; SI Fig. 1), whereas 412 individuals measured Aβ^−^ on every available PET scan (across 954 Aβ PET scans; 1-7 timepoints; Fig. 1A; SI Fig. 2). For each converter we ensured the first PET timepoint was Aβ^−^ and the last was Aβ^+^; individuals who started out Aβ^+^ but were later deemed Aβ^−^ were excluded from the Aβ PET analysis. The converter group had a maximum of 487 MRI scans available (2-14 timepoints; Fig. 1A), whereas the Aβ^−^ group had a maximum of 1850 MRI scans available (2-12 timepoints).

All participants provided written informed consent, and each study was approved by the local institutional review boards for human research. LCBC studies were approved by the Regional Ethical Committee of South-East Norway (2017/653). For BACS, ethical approval was obtained from institutional review boards at the Lawrence Berkeley National Laboratory and the University of California, Berkeley. For ADNI, ethical approval was obtained by the ADNI investigators.

### Center for Lifespan Changes in Brain and Cognition (LCBC)

#### MRI sample

1511 scans of 425 healthy individuals aged 30 to 89 years (2-7 timepoints; see SI Table 1) collected across 3 scanners (1.5T Avanto, 3T Skyra, 3T Prisma). ***PET subset:*** 242 Aβ PET scans were available for 178 individuals that had longitudinal MRI. ***General:*** Prior to participation, all individuals were screened via health and neuropsychological assessments. Generally, the LCBC sample is comprised of cognitively high-performing individuals^47,48^. The following exclusion criteria were applied across studies: evidence of neurodegenerative or neurologic disorders, conditions or injuries known to affect central nervous system (CNS) function (e.g., hypothyroidism, stroke, serious head injury), and MRI contraindications as assessed by a clinician. At baseline, participants were thoroughly screened for evidence of cognitive deficits, and excluded based on lifetime presence of psychiatric disorders and/or use of medication known to affect the CNS (e.g., benzodiazepines, antidepressants or other central nervous agents).

### Berkeley aging cohort study (BACS)

#### MRI sample

807 scans from 165 cognitively healthy, non-diagnosed individuals collected across two scanners. ***PET subset:*** 396 Aβ PET scans were available from 155 individuals (1-6 timepoints) that had longitudinal MRI. ***General:*** BACS is an ongoing longitudinal investigation of normal cognitive ageing comprised of cognitively high-performing individuals. Participants are community-dwelling adults from the Berkeley area with no history of neurological or psychiatric disorders. Eligibility criteria include a baseline Mini-Mental State Examination (MMSE) score of ≥25, normal daily functioning, no history of neurological disease, and no history of substance abuse, depression, or other psychiatric diseases. Conditions including hypertension, hyperlipidemia, and diabetes are permissible if well controlled. All participants must remain cognitively healthy while enrolled in the study.

### Alzheimer’s Disease Neuroimaging Initiative (ADNI)

#### MRI sample

2252 scans from 461 cognitively healthy, non-diagnosed individuals. We used longitudinal MRI data from individuals classed as normal controls (NC) at every diagnosed timepoint in ADNI^49^. We kept only the longitudinal MRI observations collected on the scanner field strength with the most timepoints per individual (or where equal kept the 3T scans). Hence, in the main analysis, ADNI subjects did not change field strength across time. We took this decision to help ensure the accuracy of change estimates in these legacy data, and were more at liberty to do so in ADNI due the number of longitudinal scans. Note that we identified one strong outlier in the ADNI brain change data and removed this participant from all MRI analyses (SI Fig. 26). ***PET subset***: 1046 Aβ PET scans were available from 358 cognitively healthy individuals (1-7 timepoints) that had longitudinal MRI.

### MRI acquisition and pre-processing

T1-weighted (T1w) anatomical scans from each cohort were processed using FreeSurfer’s longitudinal stream^50^ (v.7.1), yielding a reconstructed cortex for each participant and timepoint^51,52^. We extracted and modelled 70 bilateral measures of cortical thickness from the Desikan-Killiany (DK) atlas (68 regional + 2 global hemispheric measures of mean cortical thickness).

### PET analysis

Aβ positivity thresholds and centiloid calculations followed published values for BACS^28,53^, available data in ADNI (UCBERKELEY_AMY_6MM.csv downloaded from LONI on 2025-01-19), whereas we calculated the amyloid threshold in LCBC with gaussian mixture modelling and converted the values to centiloids via the GAAIN pipeline^44^ (see SI Fig. 25). LCBC PET data was acquired using [18F]flutemetamol (processing described in SI Note 1), BACS PET data was acquired using [11C]Pittsburgh compound-B (PiB; processing described in ^53^). For ADNI, PET imaging and processing followed standardized protocols, and we used both the Florbetapir (FBP) and Florbetaben (FBB) scan readouts. For each cohort, Aβ status was defined using a cortical summary SUVR that captures overall brain amyloid burden.

### Statistical analysis

For each individual in the Aβ converter group, we first calculated the time difference between their age at each MRI and age at the first Aβ^+^ scan observation. Then, we artificially truncated the longitudinal MRI data, ensuring the last MRI used to estimate brain change and structure was acquired at least X years prior to the first Aβ^+^ scan observation. For each time cutoff (X = 1-10 years before the first Aβ^+^ scan), MRI scans within that window were excluded, and we discarded those with fewer than two remaining timepoints, ensuring the data remained longitudinal. The truncated data from the converter group were then entered into mixed models.

Each time, we leveraged all of the other longitudinal MRI data to estimate the subject-specific slope and intercept of brain cortical thickness relative to a person’s age, using mixed models that benefit from incorporating as much longitudinal data as possible^37,38^. Specifically, we modelled a smooth term of age using GAMMs on the 70 cortical thickness measures (gamm4 v 0.2-6), with covariates added for sex, scanner field strength, cohort, and ICV (knots = 8). We modelled random participant effects to estimate how each person’s intercept and slope deviates from the average level and slope expected for their age. We also ran comparable GAMMs omitting the random slope term to estimate the intercept of cortical thickness, representing thickness level relative to age without estimating change (intercept-only models).

We used linear models to test group differences in the model-derived estimates. For example, to test group differences in the slopes, interpretable as the estimated degree of thickness change relative to the change expected for a person’s age, we used the slopes estimated at each cutoff as the response variable in linear models with group (converters vs Aβ^−^) as the main predictor, adjusting for mean age, sex, cohort, number of timepoints, and follow-up time (i.e., time between first and last MRI), based on the data included at each cutoff. We repeated these models with additional adjustment for quantitative amyloid levels over the years preceding amyloid positivity. For converters, we used the mean centiloid value across PET scans taken prior to the first Aβ^+^ scan; for the Aβ^−^ group, we used the mean centiloid value across all of their PET scans. We followed the same procedure to test group differences in thickness level, as estimated from the intercept-only models.

To evaluate robustness, we then ran several sensitivity analyses. To ensure the results were robust to group differences in age and longitudinal follow-up, at each cutoff we matched the Aβ^−^ group to converters based on mean age, follow-up time, and number of timepoints, using the “MatchIt” R package^54^ (SI Figs. 22-23), then re-entered the matched data from both groups into the GAMMs to estimate subject-specific effects, along with the rest of the MRI sample. Because the N of the converter group was smaller (max N = 62 [1 year cutoff]; Fig. 1d), we repeated the matching twice, drawing two one-to-one matched samples from the Aβ^−^ group (SI Figs. 22-23). This helped reduce the risk of spurious results from comparing two small groups while preserving the larger size of the Aβ^−^ group.

To rule out the possibility the group differences were driven by the field strength of the MRI scanner ^55,56^, we ran analyses including only 1.5T MRI scans. That is, we discarded all 3T scans from the data from both groups and the total combined sample, then reran the GAMMs to estimate the subject-specific random effects (MRI scans = 1848, N = 501 [1 year cutoff]), both with and without matching the groups on age and follow-up data.

To help statistically disentangle the contributions of level and change in the model-estimated estimated slopes, we used an alternative GAMM approach that modeled a tensor product interaction between each person’s mean age and time from mean age (T2 models). Specifically, we modelled how the smooth of mean age (knots = 8) varied as a function of the continuous variable time from mean age (knots = 3), correcting for sex, cohort, and ICV, with random participant intercepts and slopes. Although this enabled a cleaner separation of within-person intercept and change effects (SI Fig. 12), statistical power was reduced due to the additional model complexity and degrees of freedom involved with modelling the continuous interaction. In this model, the random slopes represent the estimated rate of cortical thickness change over time from an individual’s mean age, relative to the average change expected at their mean age. To further rule out potential confounds— particularly those relevant to modeling change—we did this using only 1.5T scans and matching the groups on age and follow-up data, and ran the models both with and without adjusting for the random intercept from our initial models. We then used the slopes estimated at each cutoff as the response variable in comparable linear models as before.

For each converter we computed predicted age at amyloid positivity (i.e., age on crossing the sample-specific Aβ^+^ threshold in centiloids; Fig. 4a), with linear models of age on the centiloid values across PET timepoints. Here, one converter was excluded as their centiloid slope over time was negative (despite their unambiguous Aβ^+^ status; SI Fig 24). Two individuals in BACS were also predicted to cross the threshold at very low ages, likely attributable to measurement error (SI Fig. 24). We then calculated the time difference between age at each MRI and age at predicted amyloid positivity, and reran the GAMMs using time-to-predicted positivity as the truncation metric (1–7 years before predicted Aβ^+^ onset). We did this using all MRI scans, 1.5T scans only, and with tensor product interaction GAMMs in 1.5T scan data. We then ran linear models as before with the slopes and intercepts estimated at each cutoff, in the complementary sample defined by predicted progression to high Aβ levels.

FDR-correction was applied within each map to adjust for multiple comparisons (70 tests), and p(FDR)<.05 was considered significant. Because statistical power varied considerably across time cutoffs and analyses, parcels that reached FDR-significance at any cutoff were considered significant at less-powered cutoffs at p<.05 (uncorrected).

Spatial correlations between the effect size maps at each cutoff and cortical Aβ deposition patterns were computed using spin tests that account for spatial autocorrelation^21^. We tested correlations with the following three Aβ maps: 1) the group-difference in SUVR between converters and the Aβ^−^ group (T-statistics from linear mixed models adjusted for tracer), 2) the group-average SUVR map across all Aβ^+^ scans (1314 scans; N = 734) in an independent ADNI sample with an MCI or AD diagnosis; 3) the group-difference in SUVR between diagnosed Aβ^+^ individuals and the Aβ^−^ group (T-statistics from linear mixed models adjusted for tracer). For regional PET analyses, the number of scans was slightly reduced due to the requirement for quality-checked regional SUVRs (rather than only global amyloid measures; converters: 260 scans, N = 76; Aβ^−^: 931 scans, N = 412). In a multiverse-style approach, we ran spatial correlations with the effect maps derived across six analyses (main and sensitivity).

To assess whether there is more thickness change in regions with higher amyloid, we ran GAMMs of time-to-predicted Aβ positivity on thickness across all MRI data in the converter group (i.e., also those acquired after conversion; 447 MRI scans; N = 76 [removing 10 MRIs from the excluded converter]. We took the average thickness across regions of high- and low Aβ accumulation (Fig. 5a). High Aβ regions were selected based on earlier analyses (Fig. 3a-d) and informed by previous reports^57^, while low Aβ regions comprised the remainder. GAMMs were adjusted for age, sex, cohort, field strength, ICV, with a random subject intercept. The difference between trajectories in high vs. low Aβ regions was tested adding a factor-smooth GAMM interaction term.

To estimate the spatiotemporal trajectory of Aβ, we ran linear mixed models of time-to-predicted Aβ positivity on the 68 regional Aβ SUVRs, adjusted for tracer, with a random subject intercept, and in the converter group only. Predicted SUVR values from these models were used to illustrate spatiotemporal Aβ accumulation across the decade leading up to Aβ positivity, and the rank order of Aβ SUVR changes was based on the point at which uptake exceeded the cerebellar gray reference region (i.e., SUVR = 1.0; Fig. 5c).

To estimate the temporal order of thickness changes, we ran GAMMs of time-to-predicted Aβ positivity on thickness, within regions consistently found to show FDR significant differences in thickness between converter and Aβ^−^ groups (and their homologues; Fig. 5d). We derived the rank order based on the point where the CI of the trajectory derivative excluded zero (i.e., where the rate-of-change first became significantly different from zero) using the “gratia” R package (v. 0.9.2). Temporal correspondence between the rank order of thickness changes and Aβ changes was tested with Pearson’s r. All statistical analyses were performed in R (v4.2.1).

## Data availability

The individual-level data supporting the results of the current study may be available upon request, given appropriate ethical and data protection approvals. Different limitations on data access apply to different samples. Participants in LCBC have not consented to share their data publicly online. Requests for the raw data can be submitted to the relevant principal investigator of each contributing study. For LCBC, data are available on request to A.M.F. For BACS, data are available on request to W.J.J. Contact details are provided in Supplementary Note 2. ADNI is available at https://adni.loni.usc.edu/data-samples/access-data/ pending application approval and compliance with the data usage agreement.

## Code availability

Code for statistical analyses will be made available at https://github.com/jamesmroe/yearsBeforeAB.

## Acknowledgements

Scripts were run on the Colossus processing cluster at the University of Oslo, and on resources provided by UNINETT Sigma2 (project NN9769K). LCBC funding: grant 302854 (FRIPRO; to Y.W.), grant 324882 (FRIPRO; to D.V-P), Peder Sather Grant Program to AMF and WJ (Peder Sather: Markers of brain activity in early Alzheimer’s Disease: The Berkeley-Oslo alliance). European Research Council under grants 283634, 725025 (to A.M.F.), and 313440 (to K.B.W.); Norwegian Research Council (to A.M.F. and K.B.W.) under grants 249931 (TOPPFORSK), The National Association for Public Health’s dementia research program, Norway (to A.M.F). The Lifebrain project is funded by the EU Horizon 2020 Grant: “Healthy minds 0–100 years: Optimising the use of European brain imaging cohorts (Lifebrain).” Grant agreement number: 732592. Some of the data used in the preparation of this article were obtained from the Alzheimer’s Disease Neuroimaging Initiative (ADNI) (National Institutes of Health Grant U01 AG024904) and DOD ADNI (Department of Defense award number W81XWH-12-2-0012). ADNI is funded by the National Institute on Aging, the National Institute of Biomedical Imaging and Bioengineering, and through generous contributions from the following: AbbVie, Alzheimer’s Association; Alzheimer’s Drug Discovery Foundation; Araclon Biotech; BioClinica, Inc.; Biogen; Bristol-Myers Squibb Company; CereSpir, Inc.; Cogstate; Eisai Inc.; Elan Pharmaceuticals, Inc.; Eli Lilly and Company; EuroImmun; F. Hoffmann-La Roche Ltd and its affiliated company Genentech, Inc.; Fujirebio; GE Healthcare; IXICO Ltd.;Janssen Alzheimer Immunotherapy Research & Development, LLC.; Johnson & Johnson Pharmaceutical Research & Development LLC.; Lumosity; Lundbeck; Merck & Co., Inc.;Meso Scale Diagnostics, LLC.; NeuroRx Research; Neurotrack Technologies; Novartis Pharmaceuticals Corporation; Pfizer Inc.; Piramal Imaging; Servier; Takeda Pharmaceutical Company; and Transition Therapeutics. The Canadian Institutes of Health Research is providing funds to support ADNI clinical sites in Canada. Private sector contributions are facilitated by the Foundation for the National Institutes of Health (www.fnih.org). The grantee organization is the Northern California Institute for Research and Education, and the study is coordinated by the Alzheimer’s Therapeutic Research Institute at the University of Southern California. ADNI data are disseminated by the Laboratory for Neuro Imaging at the University of Southern California. The ADNI researchers contributed data but did not participate in analysis or writing of this report.

