## Supplementary Information for "Cortical thickness changes precede high levels of amyloid by at least seven years"

*Roe et al.*

### SUPPLEMENTARY INFORMATION

| | ADNI NC (Obs=2252) | BACS (Obs=807) | LCBC (Obs=1511) | Overall (Obs=4570) |
|---|---|---|---|---|
| **Age** | | | | |
| Mean (SD) | 75.7 (6.45) | 78.0 (5.95) | 63.5 (14.2) | 72.1 (11.4) |
| Median [Min, Max] | 75.6 [55.8, 95.0] | 77.9 [57.4, 96.3] | 69.7 [30.1, 89.5] | 74.0 [30.1, 96.3] |
| **Sex** | | | | |
| Female | 1177 (52.3%) | 465 (57.6%) | 901 (59.6%) | 2543 (55.6%) |
| Male | 1075 (47.7%) | 342 (42.4%) | 610 (40.4%) | 2027 (44.4%) |
| **Follow-up interval** | | | | |
| Mean (SD) | 5.74 (3.13) | 7.14 (4.00) | 5.73 (4.21) | 5.99 (3.69) |
| Median [Min, Max] | 5.02 [0.507, 13.1] | 6.51 [0.504, 15.4] | 3.91 [0.522, 16.0] | 5.02 [0.504, 16.0] |

| | ADNI NC (N=461) | BACS (N=165) | LCBC (N=425) | Overall (N=1051) |
|---|---|---|---|---|
| **N Timepoints** | | | | |
| Mean (SD) | 4.89 (2.30) | 4.87 (2.62) | 3.56 (1.42) | 4.35 (2.15) |
| Median [Min, Max] | 5.00 [2.00, 11.0] | 4.00 [2.00, 14.0] | 3.00 [2.00, 7.00] | 4.00 [2.00, 14.0] |
| **Sex** | | | | |
| Female | 257 (55.7%) | 96 (58.2%) | 254 (59.8%) | 607 (57.8%) |
| Male | 204 (44.3%) | 69 (41.8%) | 171 (40.2%) | 444 (42.2%) |

**Supplementary Table 1**
Description of the total combined MRI cohort.

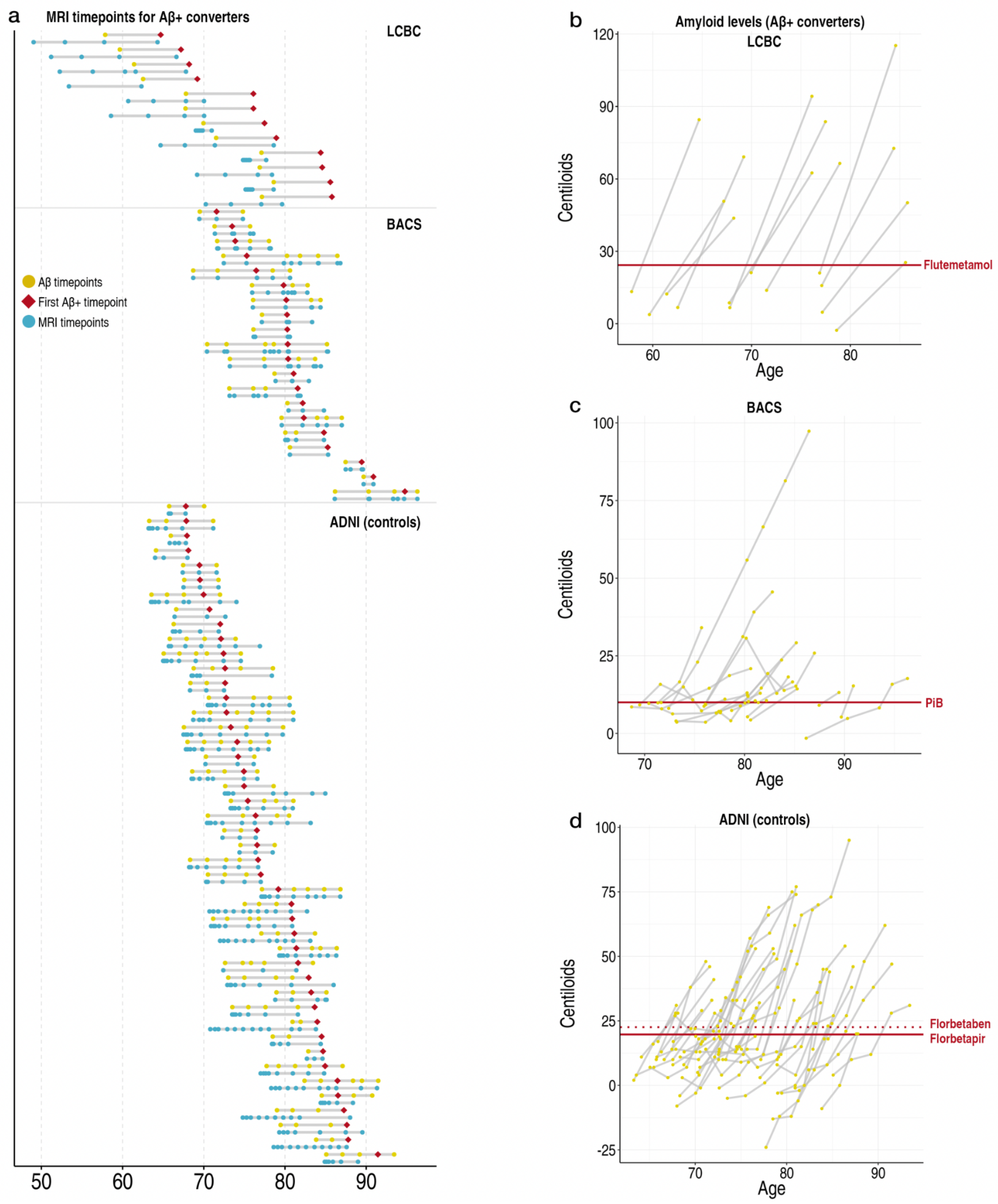

**Supplementary Figure 1**

**Amyloid PET data: Aβ+ converter group. a** Time trajectories of Aβ PET/MRI scans for the initial 77 converters (as in Fig. 1). The Aβ PET scans for each participant are in yellow and the first Aβ+ scan is depicted with a red diamond (later scans are also Aβ+). Below each is the MRI scans on that participant in blue. **b-d** Amyloid levels in centiloids for the initial 77 converters in each sample. The sample-specific threshold for amyloid positivity is shown as a solid red line. Thresholds and centiloid calculations followed published values for BACS [1,2] and ADNI (Methods), whereas we calculated the conversion threshold in LCBC with gaussian mixture modelling and converted the values to centiloids via the GAAIN pipeline[3] (SI Fig. 22).

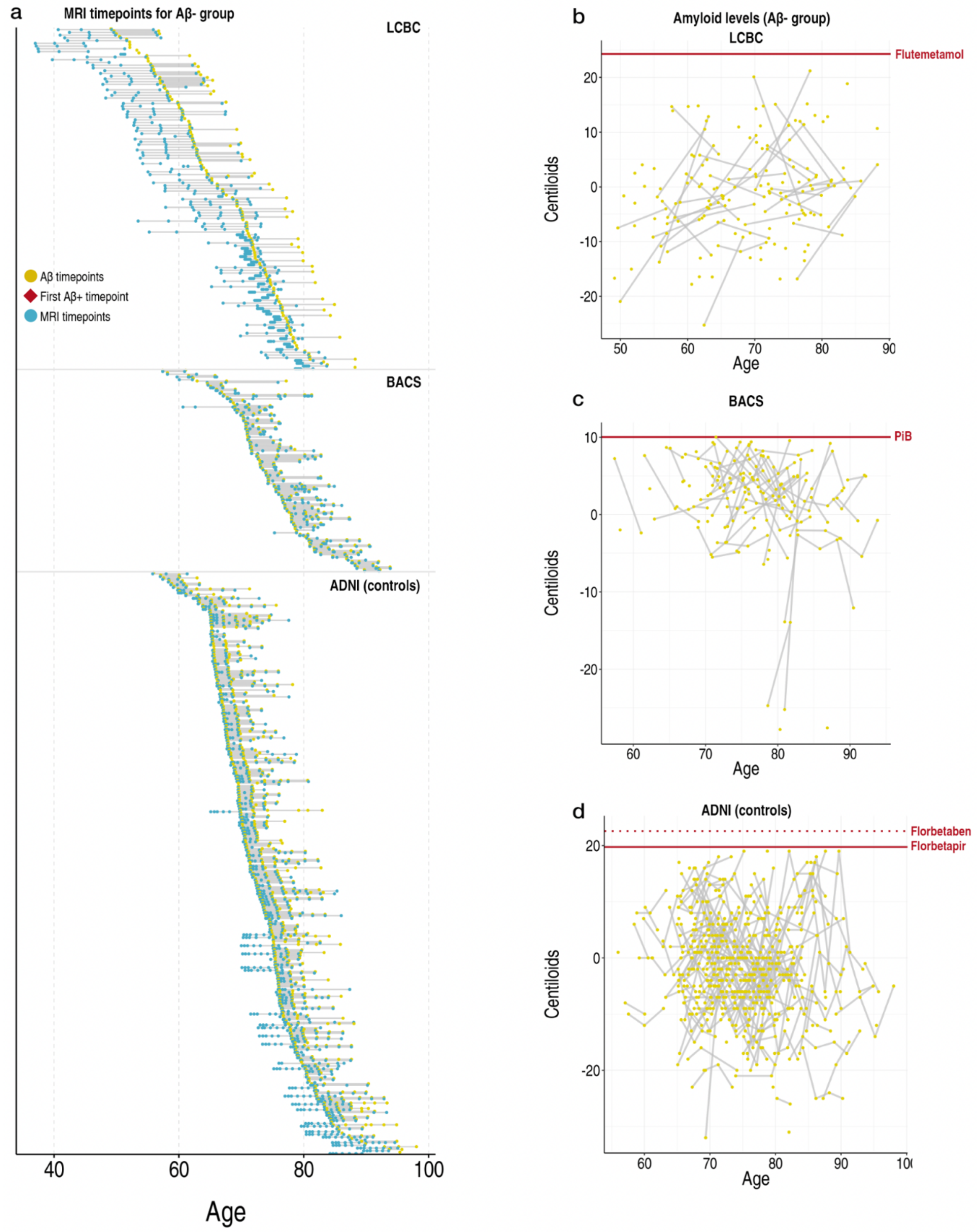

**Supplementary Figure 2**
**Amyloid PET data: Aβ- group. a** Time trajectories of Aβ PET/MRI scans for Aβ- group. The Aβ PET scans for each participant are in yellow and the first Aβ+ scan would be depicted with a red diamond (all were Aβ-). Below each is the MRI scans on that participant in blue. **b-d** Amyloid levels in centiloids for the Aβ- group in each sample. The sample-specific threshold for amyloid positivity is shown as a solid red line.

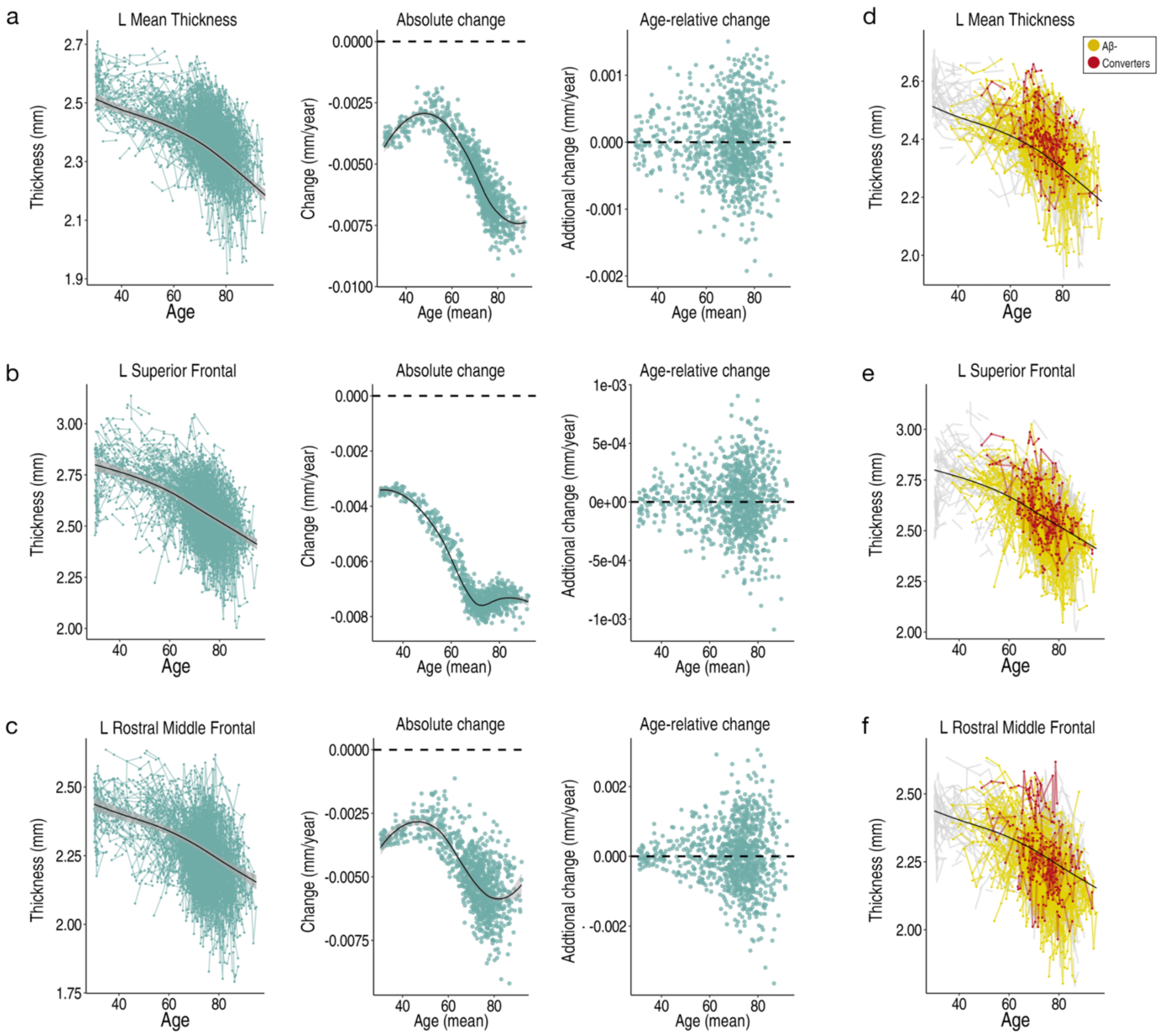

**Supplementary Figure 3**

**Cortical thickness trajectories a-c** Longitudinal data was used to estimate change in cortical thickness relative to a person's age, modelling the adult lifespan trajectories using GAMMs with random individual-specific slopes across the cortex (shown for three example thickness measures). Leftmost plots: cortical thickness age trajectories (data corrected for sex, cohort, and scanner field strength). Middle plots: estimated absolute change per individual (datapoints) as a function of their mean age across timepoints. Rightmost plots: estimated age-relative change per individual (i.e. random slopes) as a function of their mean age across timepoints. **d-f** Cortical thickness age trajectories with the data from Aβ+ converters and the Aβ- group overlain.

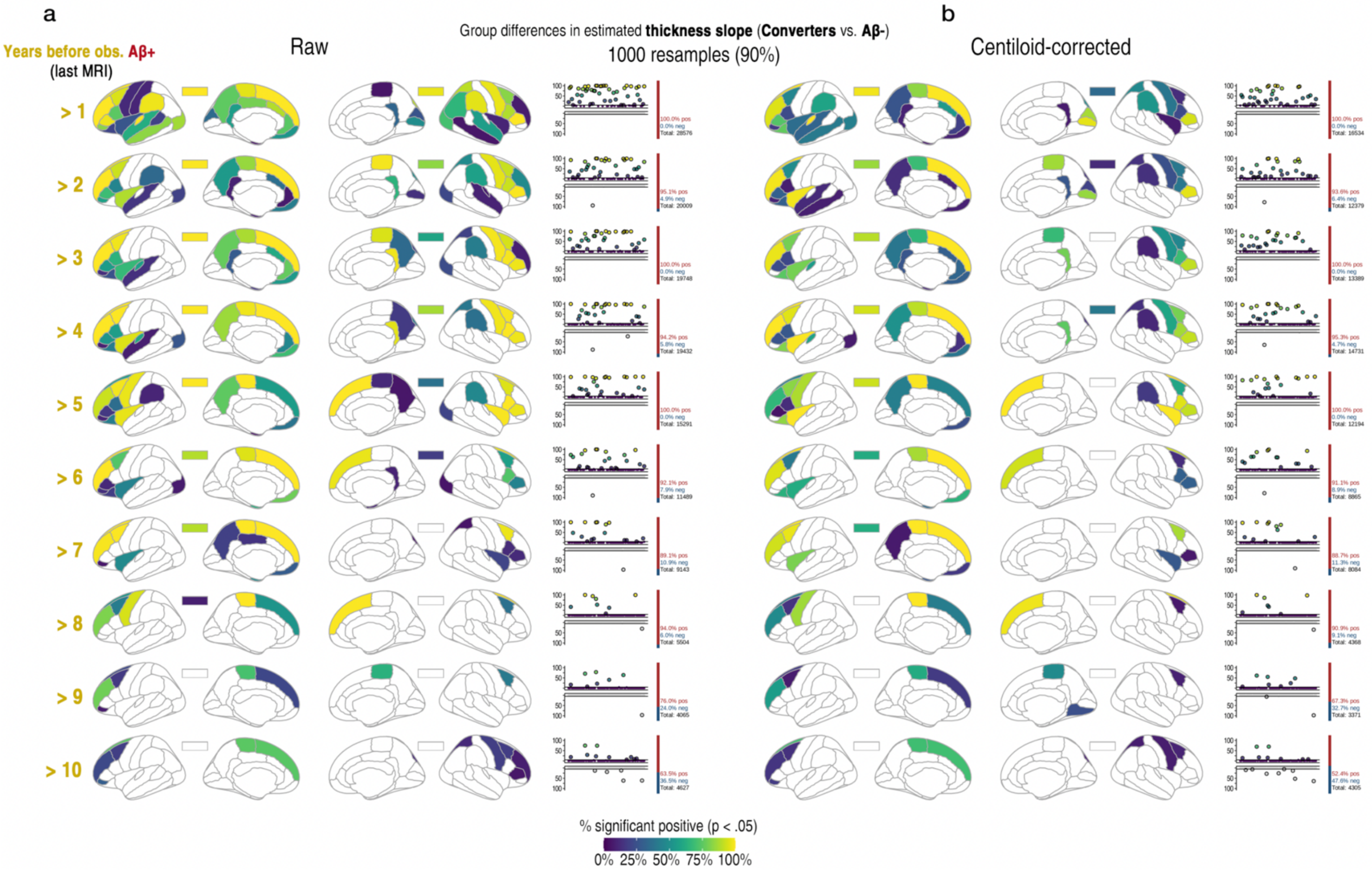

**Supplementary Figure 4**
**Resampling-based robustness check. a** Maps and plots show the number of times there was a positive significant difference at p<.05 when resampling the data drawing 90% of the estimates from each group 1,000 times (70,000 parcel-wise tests). Coloured regions exceeded the nominal 5% chance false positive rate. Each plot shows the percentage count of the number of times we observed a positive and negative effect when the difference was significant at p<.05. This illustrates the significance results were robust to sample variations and not driven by only a few observations, though caution is warranted around inferential interpretation.

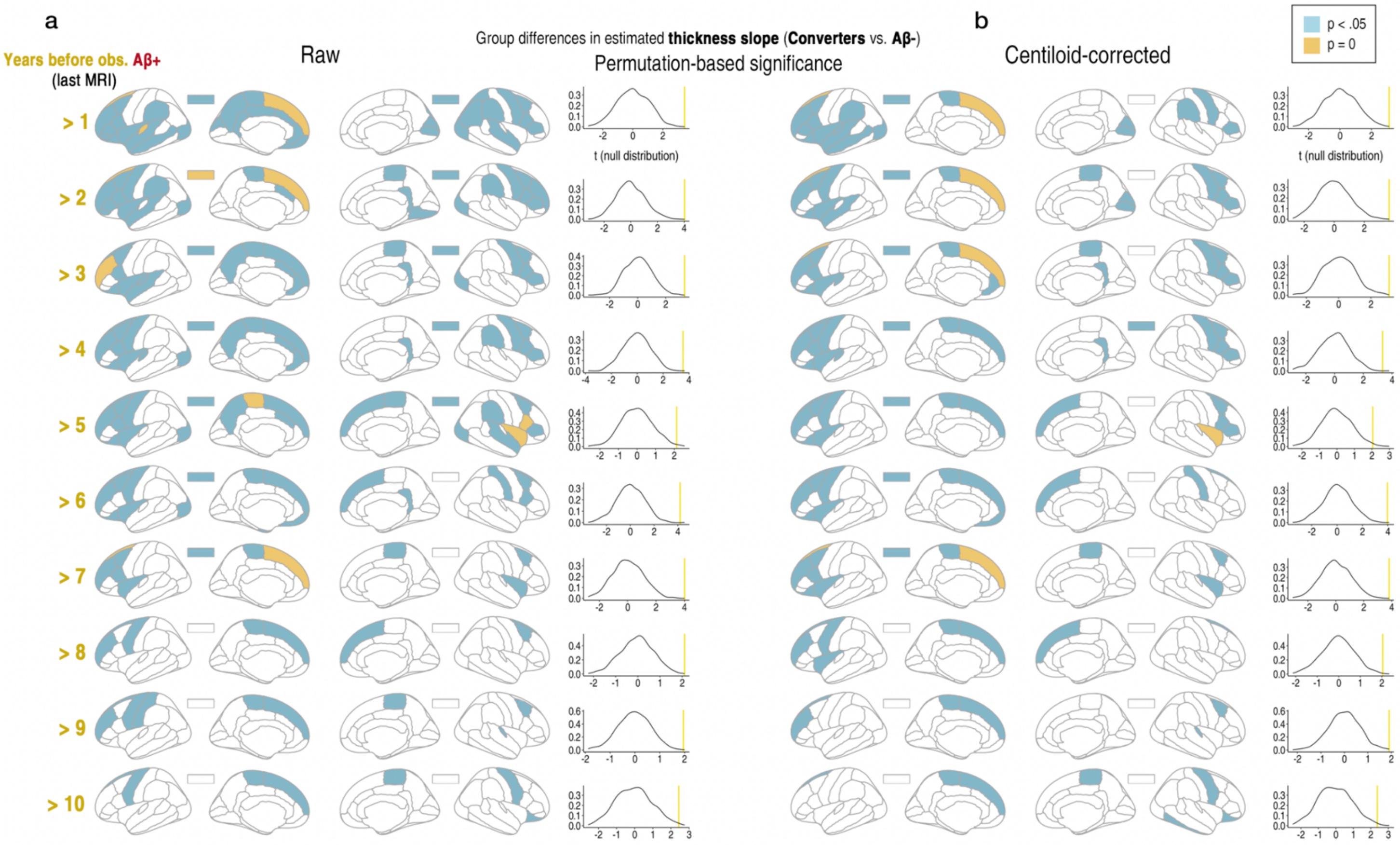

**Supplementary Figure 5**
**Permutation-based significance testing.** Significance results across 1000 permutations to evaluate whether observed group differences in estimated cortical thickness change (slopes) could be explained by group differences in variance. Null distributions were generated by randomly flipping the sign of the residuals from a null model excluding group (wild bootstrap resampling), followed by recomputing group comparisons on the resampled slopes. Any group differences arising in these null models would reflect variance imbalance alone. Blue parcels reflect empirical p-values < .05, whereas yellow parcels indicate empirical p-values of zero (i.e., the observed test statistic was more extreme than any value in the null distribution). Density plots show the observed test statistic for an example ROI (left superior frontal cortex) overlaid on the null distribution. This illustrates the observed effects were not due to group differences in the variance of the slope estimates[4].

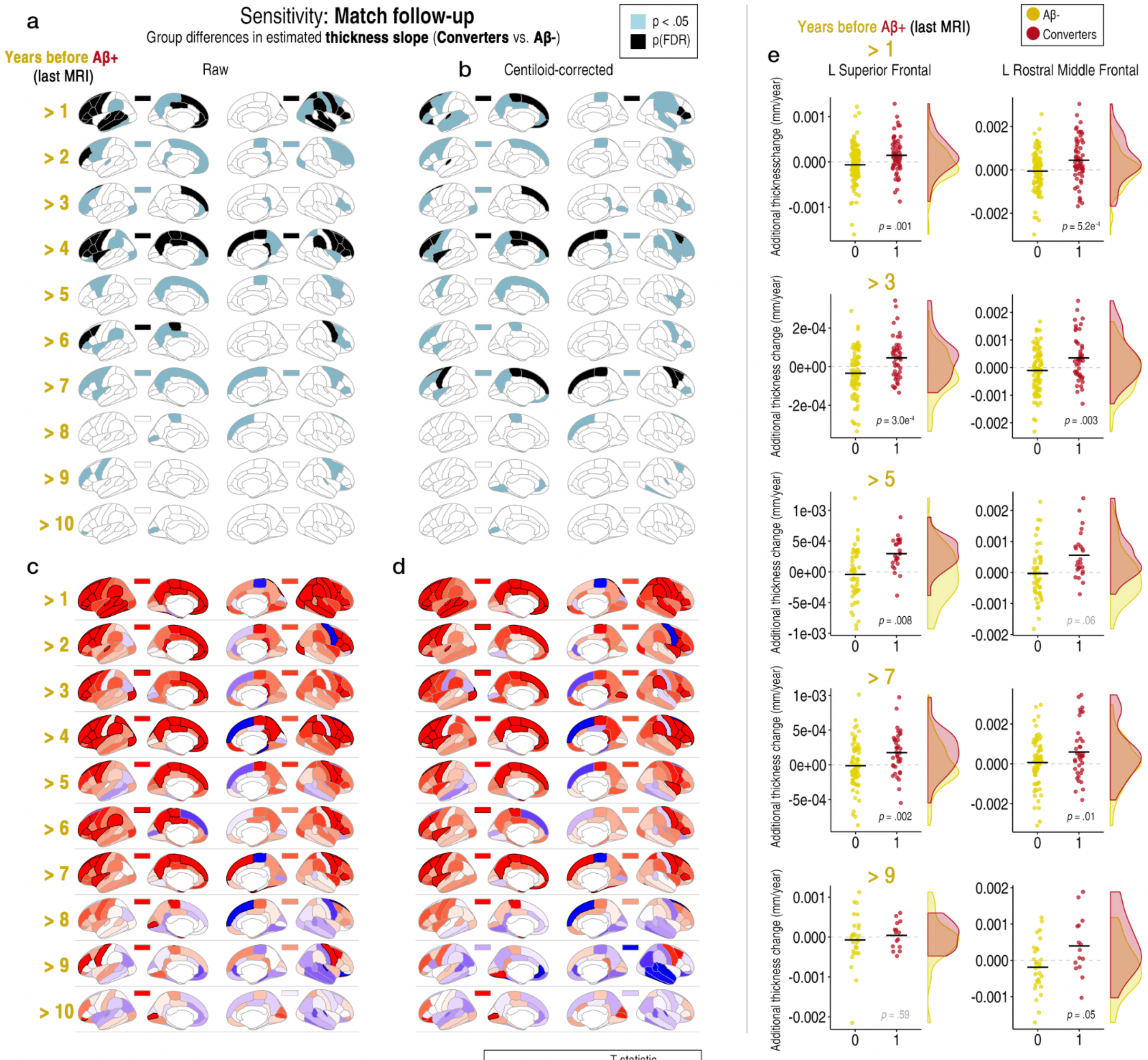

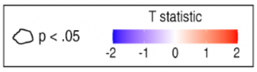

f

| Years before Aβ+ | Converters MRI Obs (N) | Aβ- MRI Obs (N) | Modelled MRI Obs (N) |
|---|---|---|---|
| > 1 | 290 (62) | 589 (124) | 3131 (749) |
| > 2 | 248 (53) | 485 (106) | 2985 (722) |
| > 3 | 214 (46) | 403 (92) | 2869 (701) |
| > 4 | 190 (42) | 359 (84) | 2801 (689) |
| > 5 | 158 (37) | 289 (74) | 2699 (674) |
| > 6 | 129 (32) | 256 (64) | 2637 (659) |
| > 7 | 93 (25) | 180 (50) | 2525 (638) |
| > 8 | 66 (19) | 129 (38) | 2447 (620) |
| > 9 | 46 (15) | 98 (30) | 2396 (608) |
| >10 | 27 (11) | 50 (22) | 2329 (596) |

**Supplementary Figure 6**

**Sensitivity analysis. Matching groups on follow-up data.** Y = model-estimated thickness slope**. a-b** Significance results after repeating the analysis drawing two one-to-one matched samples from the Aβ- group based on the number of MRI scans and time interval covered (black parcels at p[FDR] < .05; blue parcels at p < .05). **a** "raw" group differences unadjusted for quantitative amyloid. **b** Differences after adjusting for quantitative Aβ levels (mean centiloids). **c-d** T statistic maps. **e** Differences plotted for two example FDR-significant ROIs at four time cutoffs (datapoints corrected for sex, cohort, mean age, N timepoints, and interval between first and last MRI timepoint). **f** The number of MRI observations and N in the mixed-model estimation at each time cutoff, shown for both Aβ groups and the total sample submitted to the GAMMs.

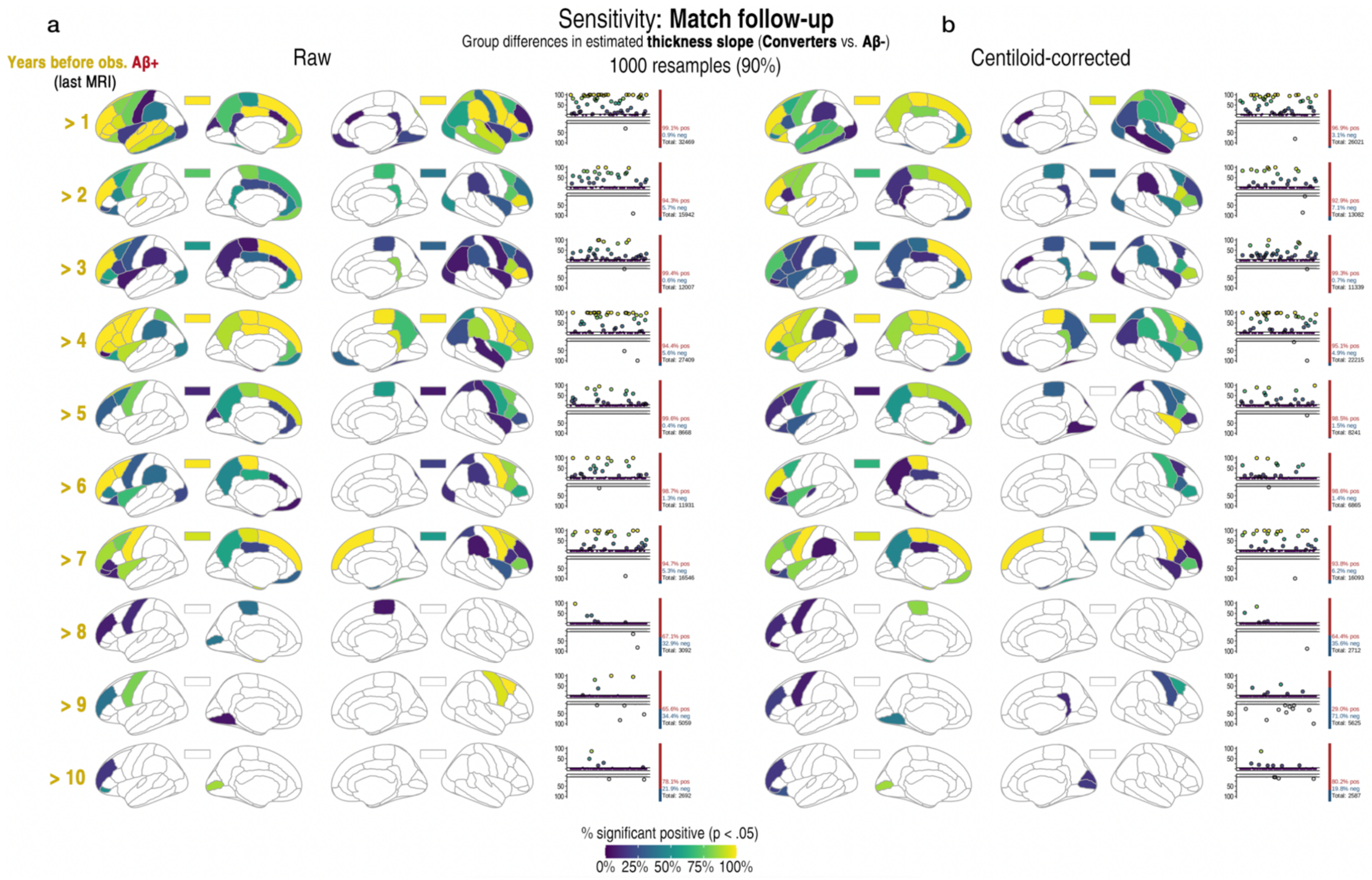

**Supplementary Figure 7**

**Resampling-based robustness check after matching groups on follow-up data. a** Maps and plots show the number of times there was a positive significant difference at p<.05 when resampling the data drawing 90% of the estimates from each group 1,000 times (70,000 parcel-wise tests). Coloured regions exceeded the nominal 5% chance false positive rate. Each plot shows the percentage count of the number of times we observed a positive and negative effect when the difference was significant at p<.05. This illustrates the reported significance results were robust to sample variations and not driven by only a few observations, though caution is warranted around inferential interpretation.

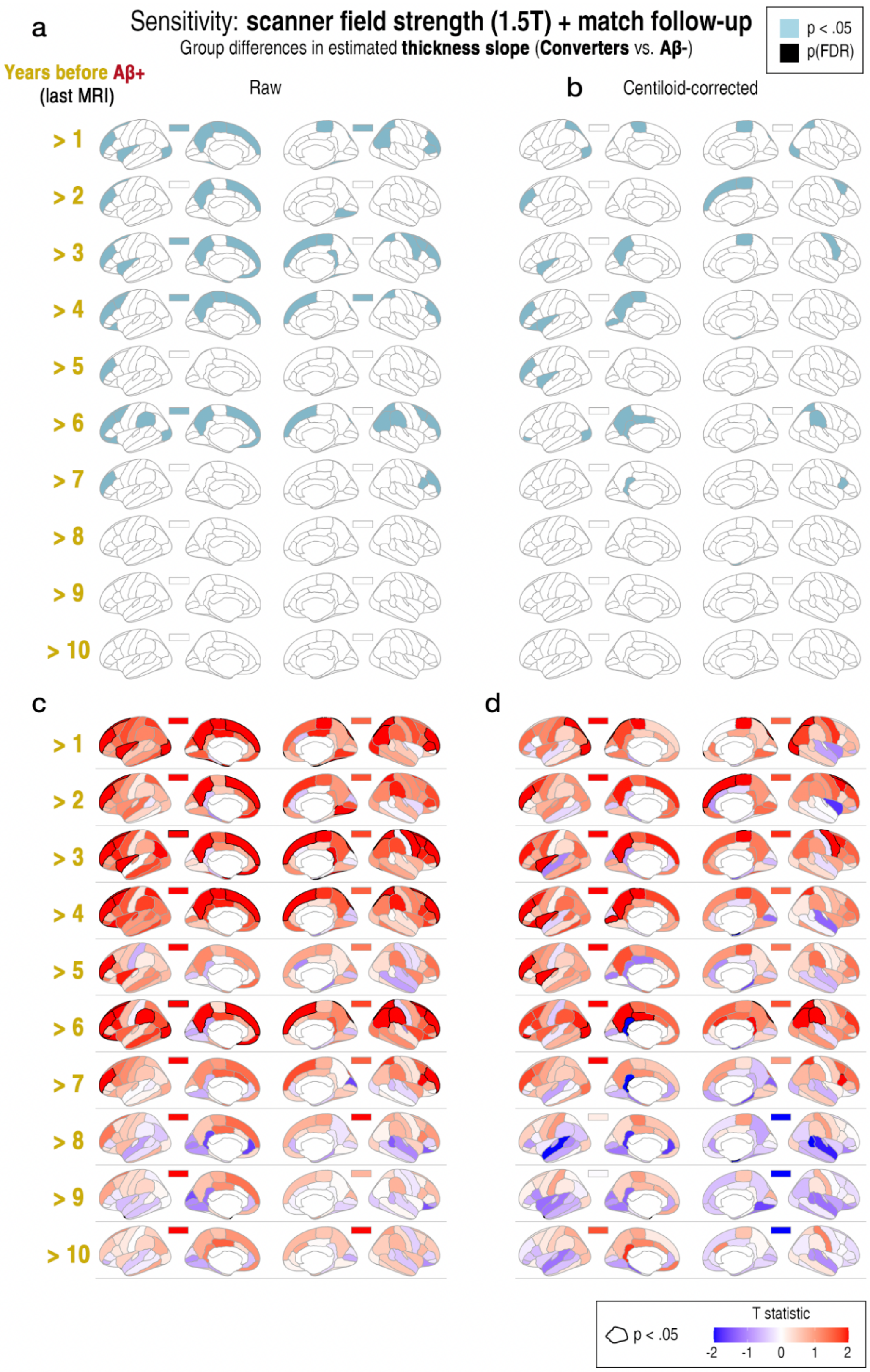

**Supplementary Figure 8**

**Sensitivity analysis: Matching groups on a single scanner field strength (1.5T) and follow-up data.** Y = model-estimated thickness slope. **a-b** Significance results after repeating the analysis drawing two one-to-one matched samples from the Aβ- group based on the number of MRI scans and time interval covered (black parcels at p[FDR] < .05; blue parcels at p < .05). **a** "raw" group differences unadjusted for quantitative amyloid. **b** Differences after adjusting for the mean centiloid values across PET scans prior to the first Aβ$^{+}$ scan. **c-d** T statistic maps.

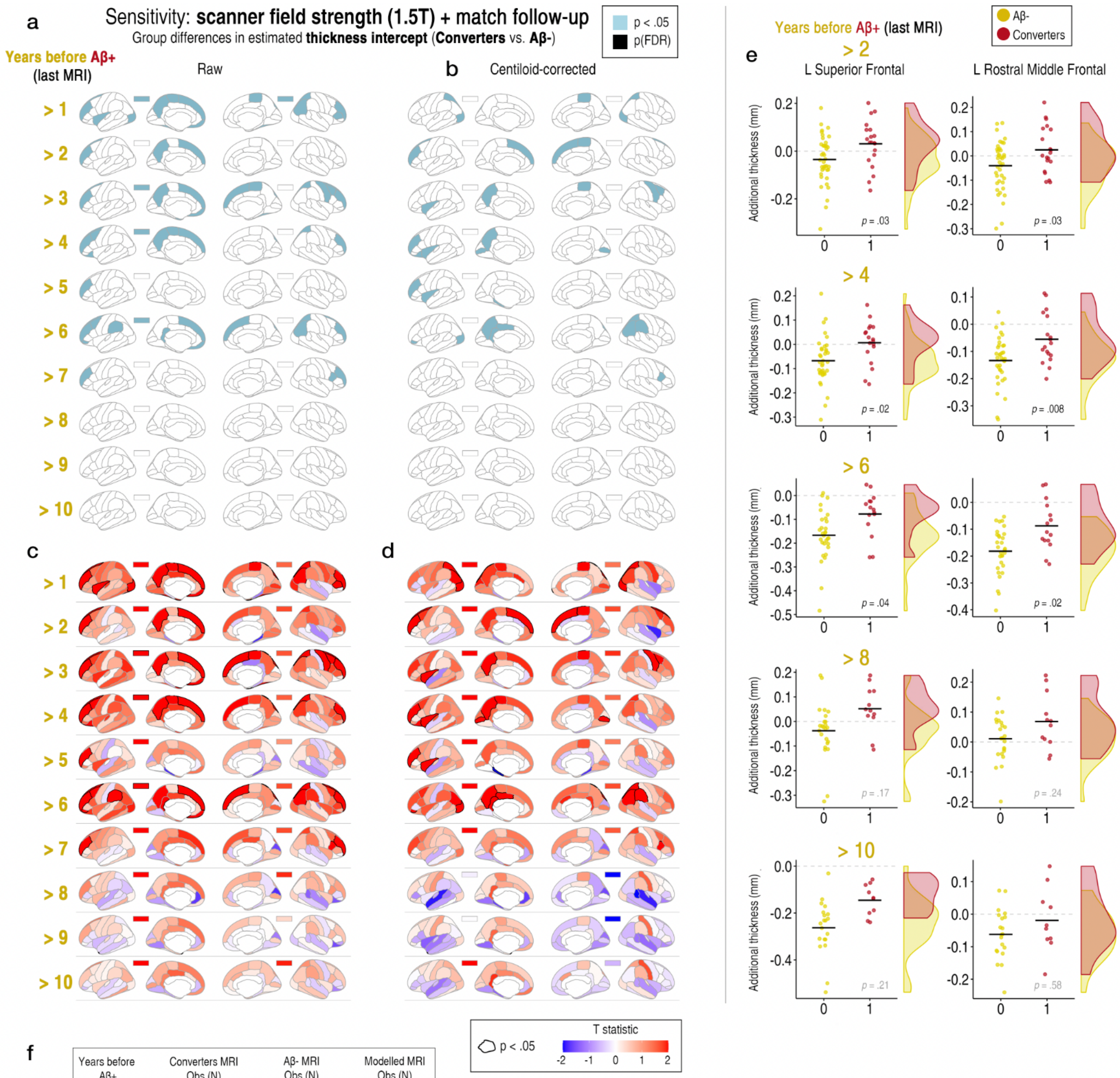

f

| Years before Aβ+ | Converters MRI Obs (N) | Aβ- MRI Obs (N) | Modelled MRI Obs (N) |
|---|---|---|---|
| > 1 | 91 (22) | 170 (44) | 1426 (388) |
| > 2 | 82 (20) | 159 (40) | 1406 (382) |
| > 3 | 76 (19) | 162 (38) | 1403 (379) |
| > 4 | 71 (18) | 161 (36) | 1397 (376) |
| > 5 | 64 (16) | 127 (32) | 1356 (370) |
| > 6 | 57 (15) | 128 (30) | 1350 (367) |
| > 7 | 48 (14) | 99 (28) | 1312 (364) |
| > 8 | 36 (12) | 70 (24) | 1271 (358) |
| > 9 | 26 (10) | 52 (20) | 1243 (352) |
| >10 | 24 (10) | 45 (20) | 1234 (352) |

**Supplementary Figure 9**

**Sensitivity analysis: Matching groups on a single scanner field strength (1.5T) and follow-up data**. Y = model-estimated thickness intercept (intercept-only random effect model; Methods). **a-b** Significance results after repeating the analysis drawing two one-to-one matched samples from the Aβ- group based on the number of MRI scans and time interval covered (black parcels at p[FDR] < .05; blue parcels at p < .05). **a** "raw" group differences unadjusted for quantitative amyloid. **b** Differences after adjusting for the mean centiloid values across PET scans prior to the first Aβ$^+$ scan. **c-d** T statistic maps. **e** Differences plotted for two example ROIs at four time cutoffs. Datapoints corrected for sex, cohort, mean age, N timepoints, and interval between first and last MRI timepoint. **f** The number of MRI observations and N in the mixed-model estimation at each time cutoff, shown for both Aβ groups and the total sample submitted to the GAMMs

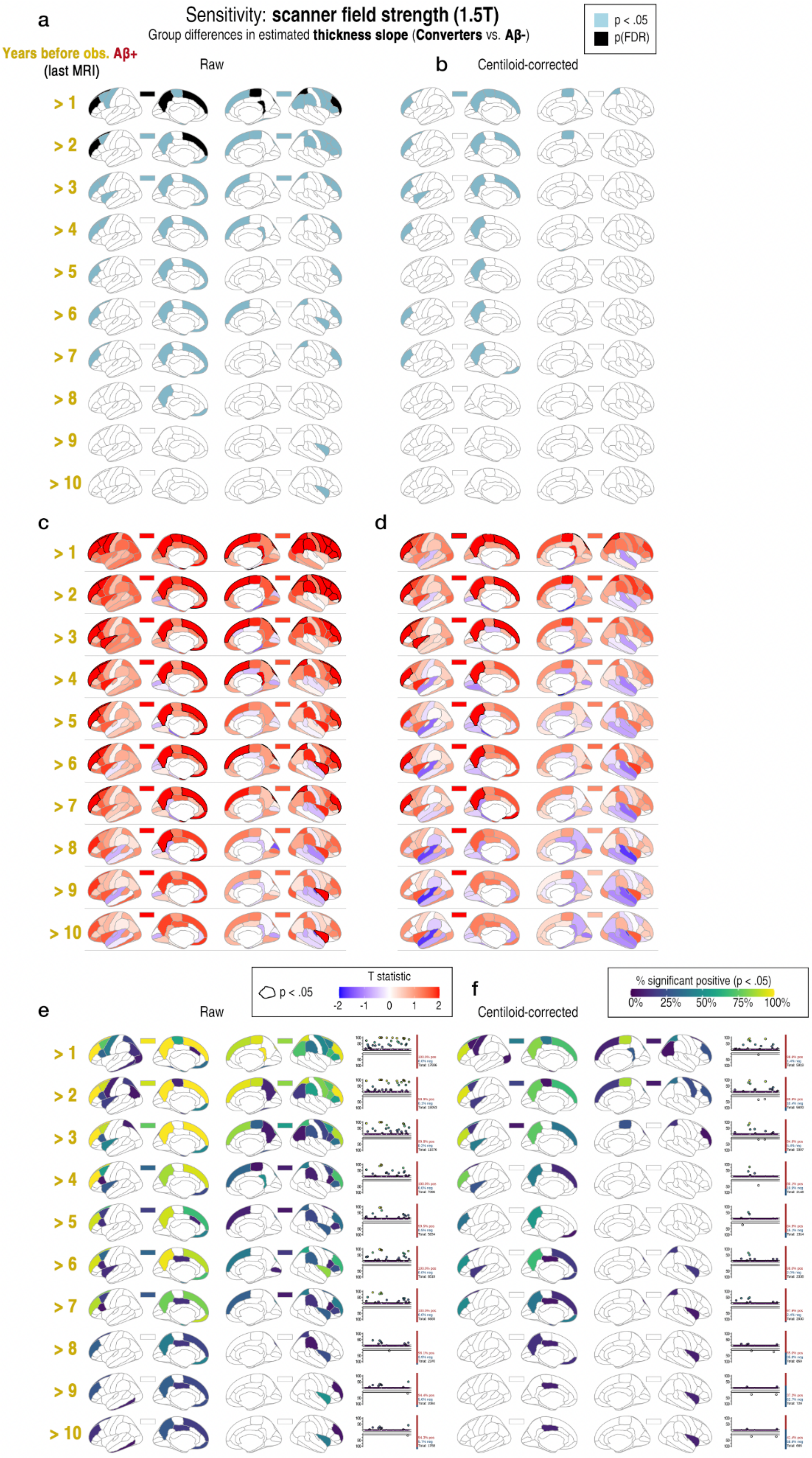

a
Sensitivity: scanner field strength (1.5T)
Group differences in estimated thickness slope (Converters vs. Aβ-)
p < .05
p(FDR)
Years before obs. Aβ+ (last MRI)
Raw
b
Centiloid-corrected
> 1
> 2
> 3
> 4
> 5
> 6
> 7
> 8
> 9
> 10
c
d
T statistic
p < .05
-2 -1 0 1 2
e
Raw
f
Centiloid-corrected
% significant positive (p < .05)
0% 25% 50% 75% 100%

**Supplementary Figure 10**

**Sensitivity analysis: Matching groups on a single scanner field strength only (1.5T).** Y = model-estimated thickness slope. **a-b** Significance results after repeating the analysis using only 1.5T scans (black parcels at p[FDR] < .05; blue parcels at p < .05). **a** "raw" group differences unadjusted for quantitative amyloid. **b** Differences after adjusting for the mean centiloid values across PET scans prior to the first $A\beta^{+}$ scan. **c-d** T statistic maps. **e-f** Resampling-based robustness check. Maps and plots show the number of times there was a positive significant difference at p<.05 when resampling the data drawing 90% of the estimates from each group 1,000 times (70,000 parcel-wise tests). Coloured regions exceeded the nominal 5% chance false positive rate. Each plot shows the percentage count of the number of times we observed a positive and negative effect when the difference was significant at p<.05. This illustrates the reported significance results were robust to sample variations and not driven by only a few observations, though caution is warranted around inferential interpretation.

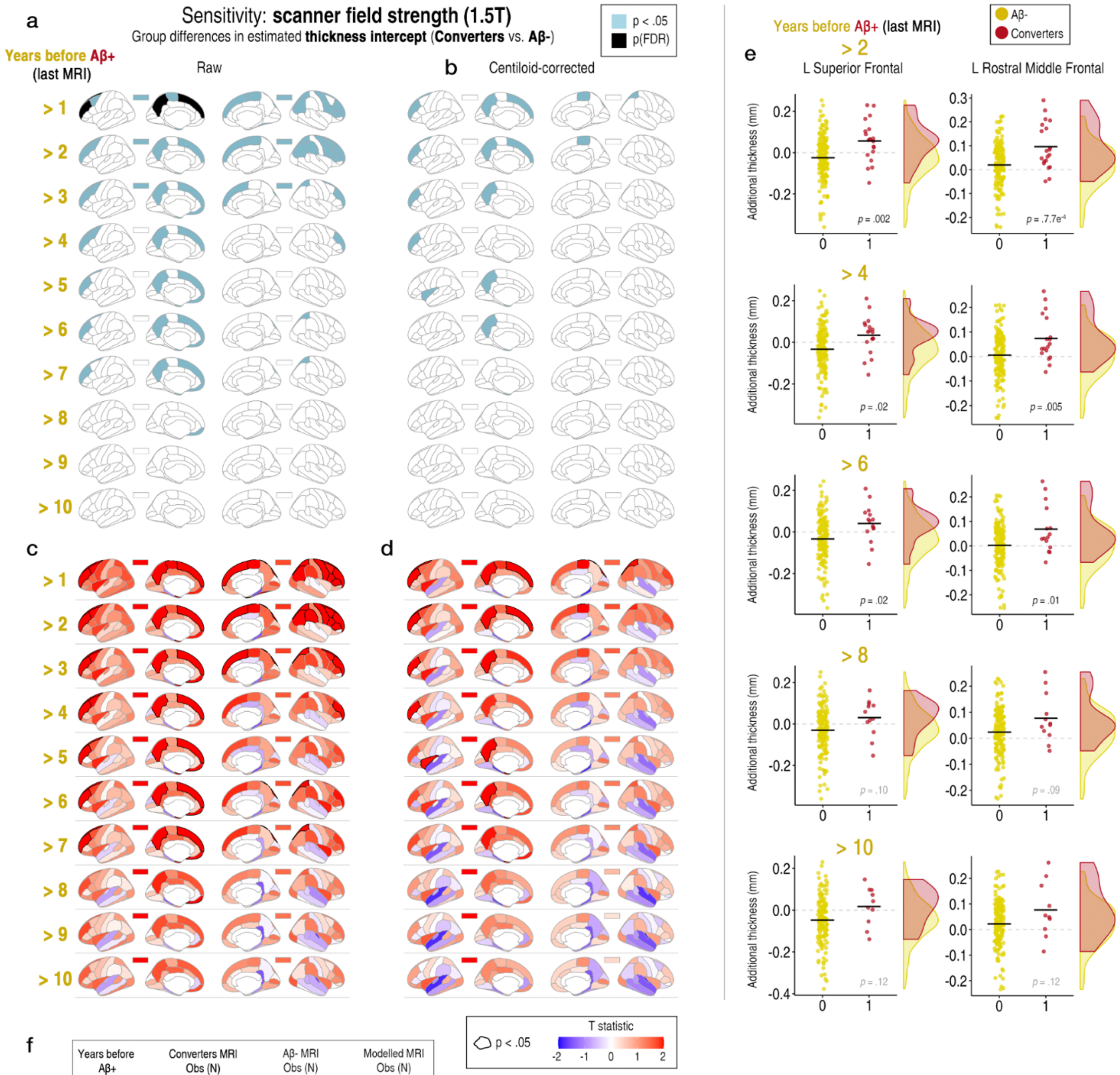

f

| Years before Aβ+ | Converters MRI Obs (N) | Aβ- MRI Obs (N) | Modelled MRI Obs (N) |
|---|---|---|---|
| > 1 | 91 (22) | 592 (157) | 1848 (501) |
| > 2 | 82 (20) | 592 (157) | 1839 (499) |
| > 3 | 76 (19) | 592 (157) | 1833 (498) |
| > 4 | 71 (18) | 592 (157) | 1828 (497) |
| > 5 | 64 (16) | 592 (157) | 1821 (495) |
| > 6 | 57 (15) | 592 (157) | 1814 (494) |
| > 7 | 48 (14) | 592 (157) | 1805 (493) |
| > 8 | 36 (12) | 592 (157) | 1793 (491) |
| > 9 | 26 (10) | 592 (157) | 1783 (489) |
| >10 | 24 (10) | 592 (157) | 1781 (489) |

**Supplementary Figure 11**
**Sensitivity analysis: Matching groups on a single scanner field strength only (1.5T).** Y = model-estimated thickness intercept (intercept-only random effect models; Methods). **a-b** Significance results after repeating the analysis using only 1.5T scans (black parcels at p[FDR] < .05; blue parcels at p < .05). **a** "raw" group differences unadjusted for quantitative amyloid. **b** Differences after adjusting for the mean centiloid values across PET scans prior to the first Aβ$^{+}$ scan. **c-d** T statistic maps. **e** Differences plotted for two example FDR-significant ROIs at four time cutoffs. Datapoints corrected for sex, cohort, mean age, N timepoints, and interval between first and last MRI timepoint. **f** The number of MRI observations and N in the mixed-model estimation at each time cutoff, shown for both Aβ groups and the total sample submitted to the GAMMs.

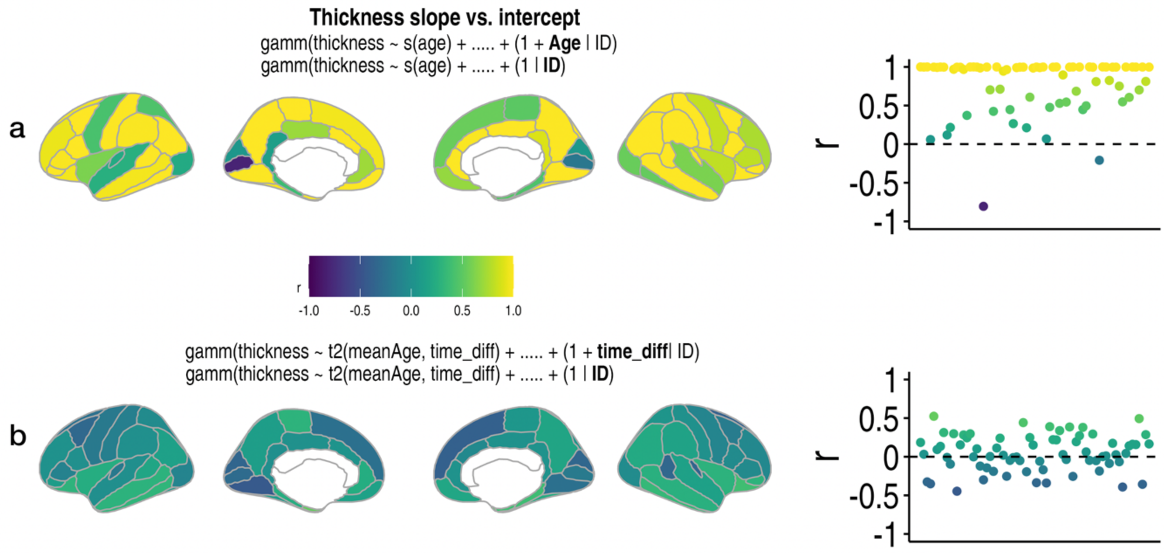

**Supplementary Figure 12**
Correlations between the model-estimated random slope and intercept (intercept-only random effects model) in our initial models. To statistically help statistically separate the effects, we ran alternative models using a tensor smooth interaction between mean age and time from mean age, and used the slopes from these models to test group differences in cortical thickness change, with (Fig. 13) and without (SI Fig. 12) correcting for the intercept from the intercept-only model.

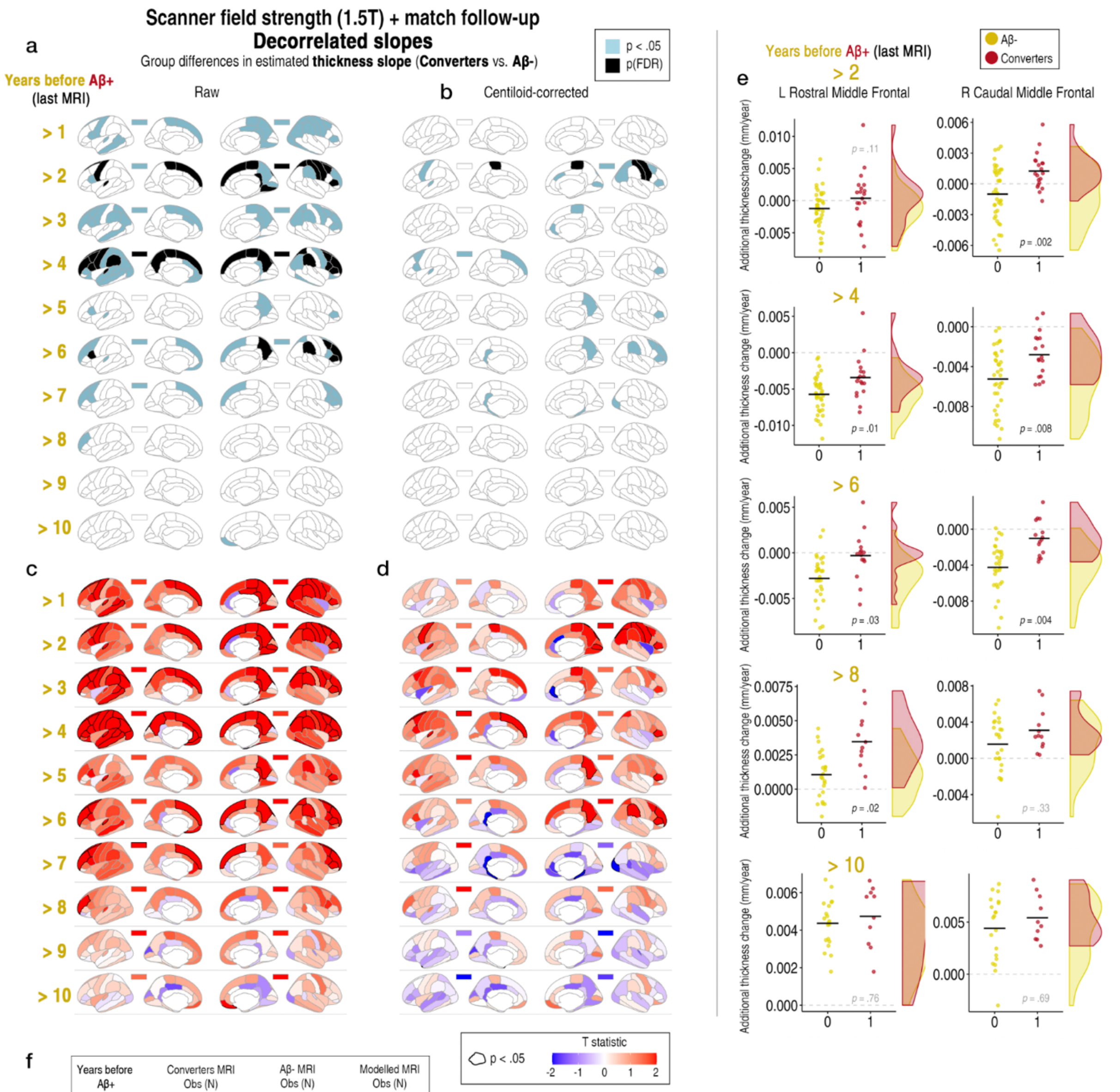

f

| Years before Aβ+ | Converters MRI Obs (N) | Aβ- MRI Obs (N) | Modelled MRI Obs (N) |
|---|---|---|---|
| > 1 | 91 (22) | 170 (44) | 1426 (388) |
| > 2 | 82 (20) | 159 (40) | 1406 (382) |
| > 3 | 76 (19) | 162 (38) | 1403 (379) |
| > 4 | 71 (18) | 161 (36) | 1397 (376) |
| > 5 | 64 (16) | 127 (32) | 1356 (370) |
| > 6 | 57 (15) | 128 (30) | 1350 (367) |
| > 7 | 48 (14) | 99 (28) | 1312 (364) |
| > 8 | 36 (12) | 70 (24) | 1271 (358) |
| > 9 | 26 (10) | 52 (20) | 1243 (352) |
| > 10 | 24 (10) | 45 (20) | 1234 (352) |

**Supplementary Figure 13**

**Differences in cortical thickness change between matched Aβ+ converter and Aβ- groups (models isolating slope from intercept effects). a-b** Significant differences in the slope of cortical thickness between Aβ+ converters and Aβ- groups matched on scanner field strength (1.5T scans only), age and follow-up data, using alternative GAMMs to statistically separate slope from intercept effects (tensor smooth interaction models; Methods). Models shown are unadjusted for the intercept (compare with SI Fig. 13 for adjusted results). Black parcels depict FDR-significant hits, whereas blue depicts significance at p<.05 (uncorrected). The rectangles above each brain hemisphere depict the results for mean cortical thickness. **a** Depicts "raw" differences uncorrected for amyloid levels, whereas **b** depicts differences after correcting for quantitative amyloid (mean centiloids across PET scans prior to the first $A\beta^{+}$ scan). **c-d** Corresponding T maps (unthresholded). **e** Differences plotted for two example FDR-significant ROIs at four time cutoffs. Datapoints corrected for sex, cohort, mean age, N timepoints, and interval between first and last MRI timepoint. **f** The number of MRI observations and N in the mixed-model estimation at each time cutoff, shown for both Aβ groups and the total sample submitted to the GAMMs.

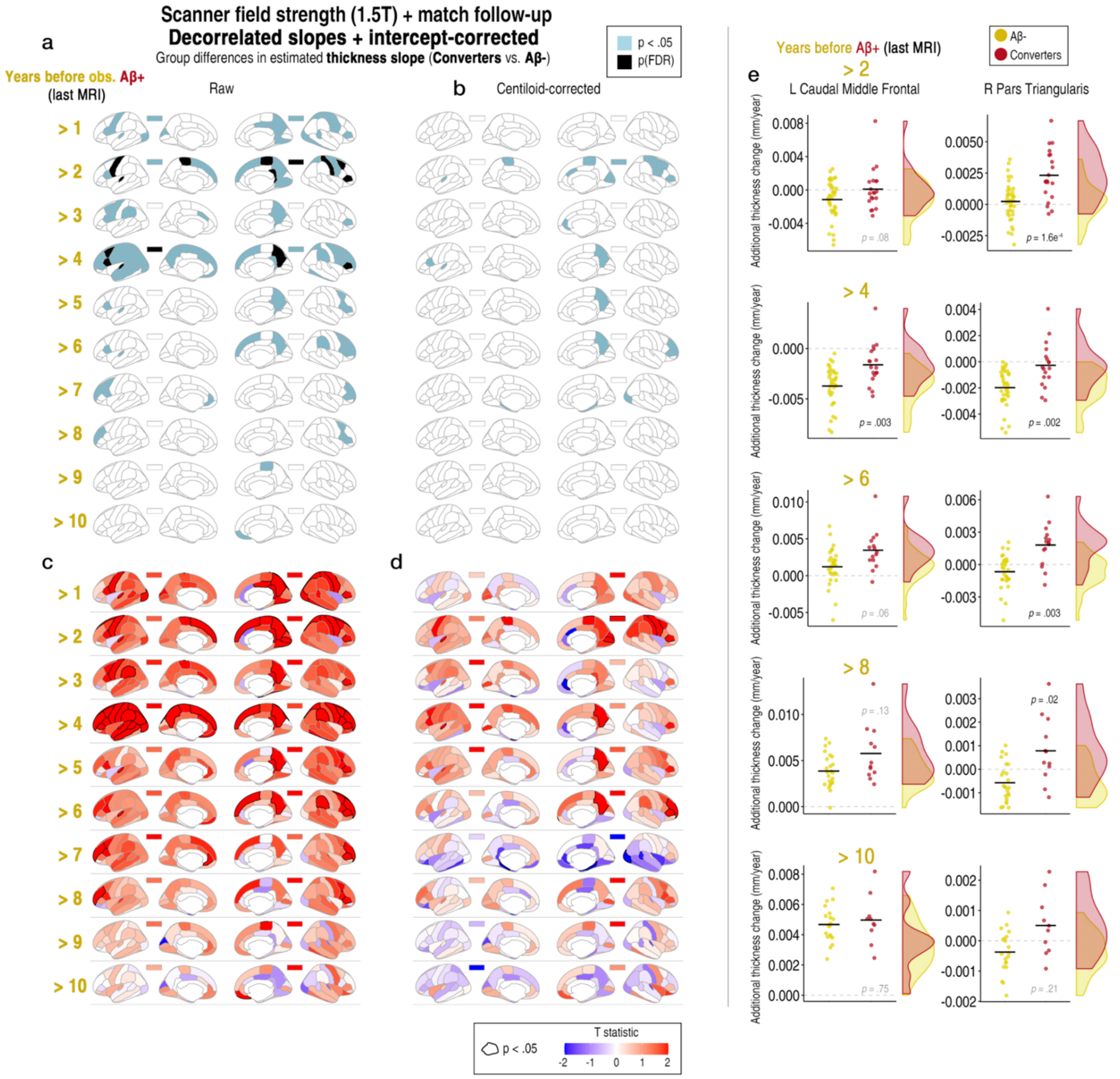

**Supplementary Figure 14**

**Differences in cortical thickness change between matched Aβ+ converter and Aβ- groups (models isolating slope from intercept effects). a-b** Significant differences in cortical thickness change between Aβ+ converters and Aβ- groups matched on scanner field strength (1.5T scans only), age and follow-up data, using alternative GAMMs to statistically separate slope from intercept effects (tensor smooth interaction models; Methods). Models shown are additionally adjusted for the intercept (intercept-only random effect models). Black parcels depict FDR-significant hits, whereas blue depicts significance at p<.05 (uncorrected). The rectangles above each brain hemisphere depict the results for mean cortical thickness. **a** Depicts "raw" differences uncorrected for amyloid levels, whereas **b** depicts differences after correcting for quantitative amyloid (mean centiloids across PET scans prior to the first Aβ$^{+}$ scan). **c-d** Corresponding T maps (unthresholded). **e** Differences plotted for two example FDR-significant ROIs at four time cutoffs. Datapoints corrected for sex, cohort, mean age, N timepoints, and interval between first and last MRI timepoint. **f** The number of MRI observations and N in the mixed-model estimation at each time cutoff, shown for both Aβ groups and the total sample submitted to the GAMMs.

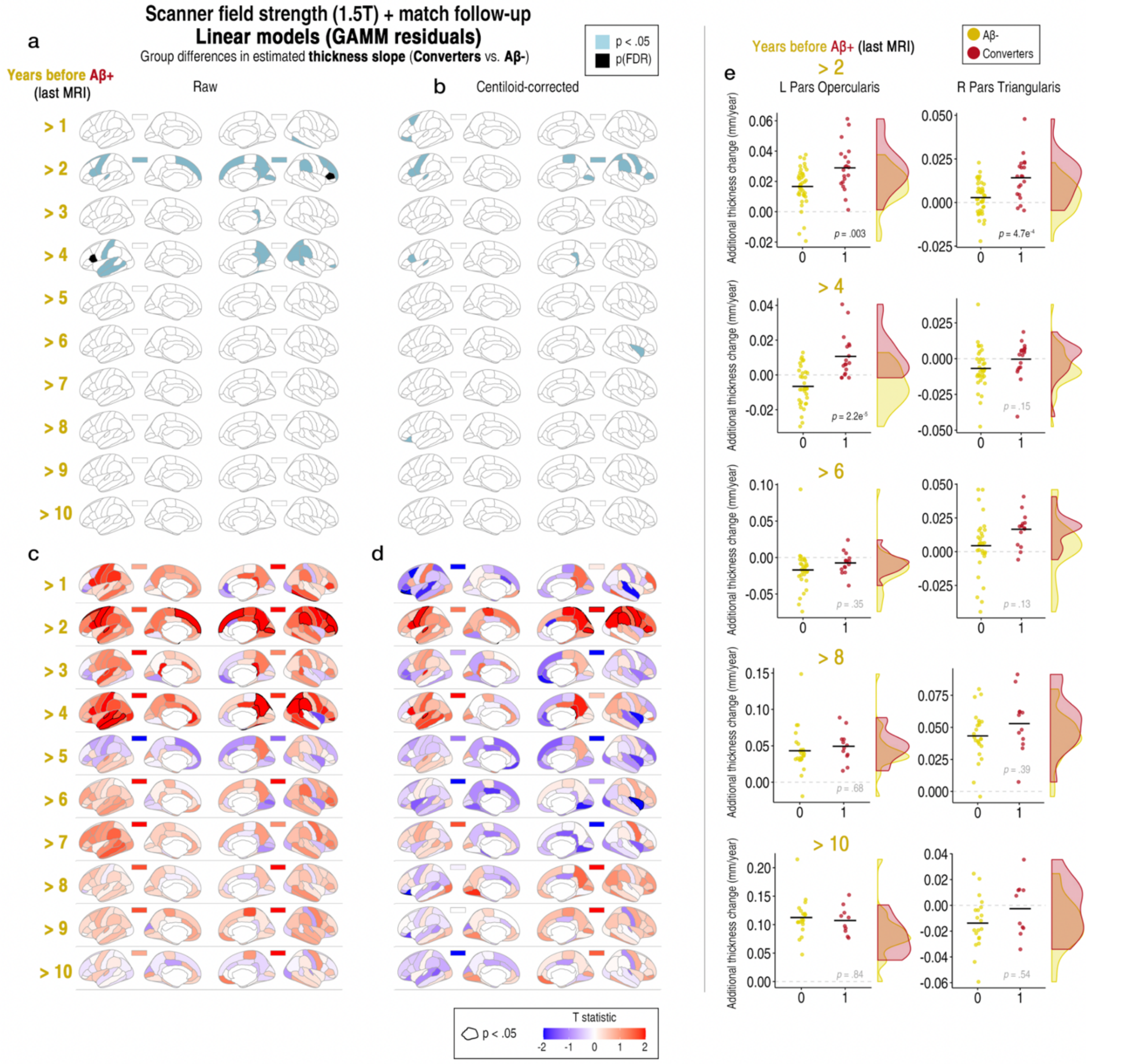

**Supplementary Figure 15**

**Matching groups on a single scanner field strength (1.5T) and follow-up data (simple linear change)**. Y = linear slope across timepoints after regressing out nonlinear age (GAMM residuals [intercept-only random effect models], i.e. age-relative thickness change computed "manually"). Note that as this analysis involved computing simple linear slopes across progressively truncated data (see Table in SI Fig. 10E), we removed outliers >5SD prior to performing the group-difference tests. In contrast, the mixed-effects approach will be less influenced by outliers due to the shrinkage effect. This limits the influence of extreme data points by estimating random effects from a probability distribution, the parameters of which are derived from the data. As more longitudinal measures are incorporated, the distribution becomes more robust, reducing the influence of extreme slopes and pulling them closer to the mean[5,6] **a** Depicts "raw" differences uncorrected for amyloid levels, whereas **b** depicts differences after correcting for quantitative amyloid (mean centiloids across PET scans prior to the first $A\beta^+$ scan). **c-d** Corresponding T maps (unthresholded). **e** Differences plotted for two example FDR-significant ROIs at four time cutoffs. Datapoints corrected for sex, cohort, mean age, N timepoints, and interval between first and last MRI timepoint.

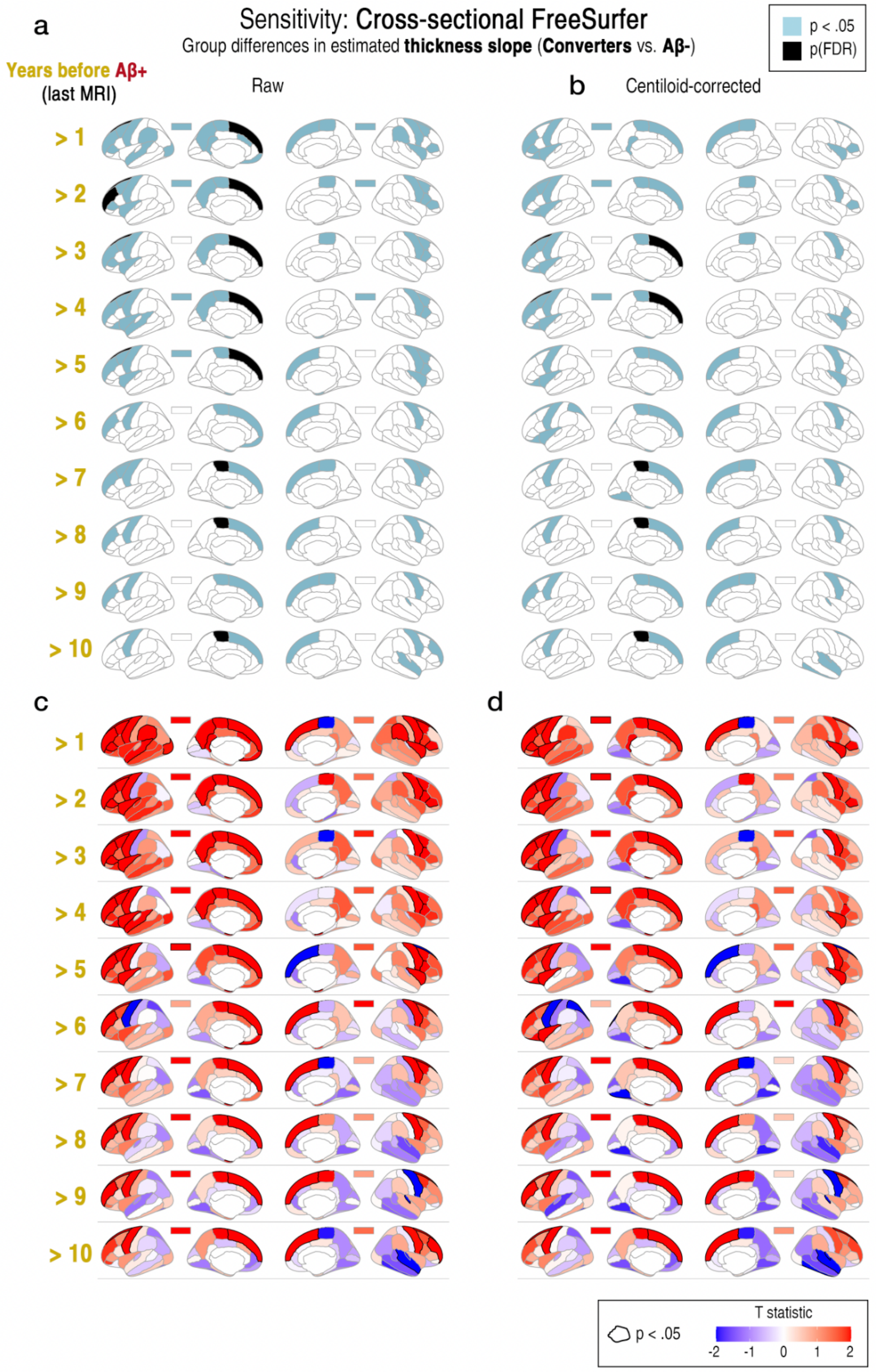

**Supplementary Figure 16**
Results from the main analysis (Fig. 1) using cross-sectionally processed MRI scans with FreeSurfer.

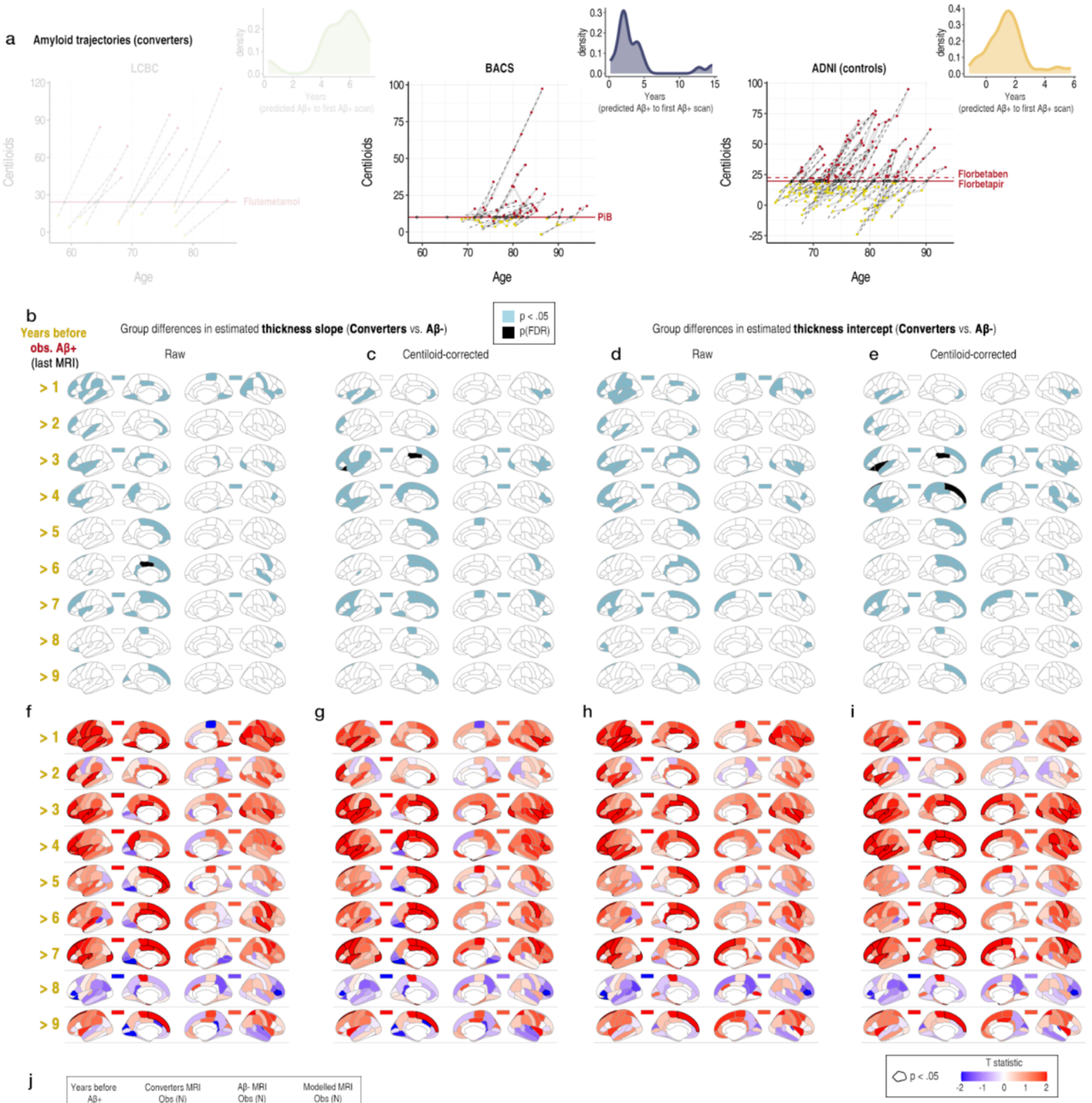

j

| Years before Aβ+ | Converters MRI Obs (N) | Aβ- MRI Obs (N) | Modelled MRI Obs (N) |
|---|---|---|---|
| > 1 | 241 (50) | 461 (100) | 2954 (713) |
| > 2 | 199 (41) | 370 (82) | 2821 (686) |
| > 3 | 165 (34) | 312 (68) | 2729 (665) |
| > 4 | 141 (30) | 294 (60) | 2687 (653) |
| > 5 | 109 (25) | 219 (50) | 2580 (638) |
| > 6 | 80 (20) | 156 (40) | 2488 (623) |
| > 7 | 55 (14) | 114 (28) | 2421 (605) |
| > 8 | 34 (8) | 63 (16) | 2349 (587) |
| > 9 | 22 (5) | 43 (10) | 2317 (578) |

**Supplementary Figure 17**
**Differences in estimated slope and intercept of cortical thickness between matched converter and Aβ- groups after discarding the data from LCBC converters (GAMMs). a-c** Amyloid trajectories in centiloids for the Aβ+ converters in each sample (linear model across PET timepoints; raw data in SI Figs. 1-2; see also SI Fig. 13). The sample-specific threshold for Aβ positivity is shown as a solid red line (Methods), and predicted age on crossing the threshold is shown by a white dot (note we used the Florbetapir threshold in ADNI which was lower). Inset plots show the time from predicted to observed Aβ positivity (i.e. first Aβ+ scan) for each sample. Note that as LCBC only had two PET timepoints, more individuals were predicted to cross the Aβ+ threshold several years before they were observed Aβ+, illustrating how time cutoffs based on years to the first Aβ+ scan should be interpreted with caution. We therefore reran the analysis discarding the data from LCBC converters in the GAMMs. **b-c** Significant differences in the model-estimated slope of cortical thickness, with and without adjusting for amyloid levels (mean centiloids across PET scans prior to the first Aβ$^{+}$ scan). **d-e** Significant differences in the model-estimated intercept of cortical

thickness, with and without adjusting for amyloid. Black parcels depict FDR-significant hits, whereas blue depicts significance at p<.05 (uncorrected). The rectangles above each brain hemisphere depict the results for mean cortical thickness. **f-i** Corresponding T maps (unthresholded). **i f** The number of MRI observations and N in the mixed-model estimation at each time cutoff, shown for both Aβ groups and the total sample submitted to the GAMMs. Groups were matched on age and follow-up data.

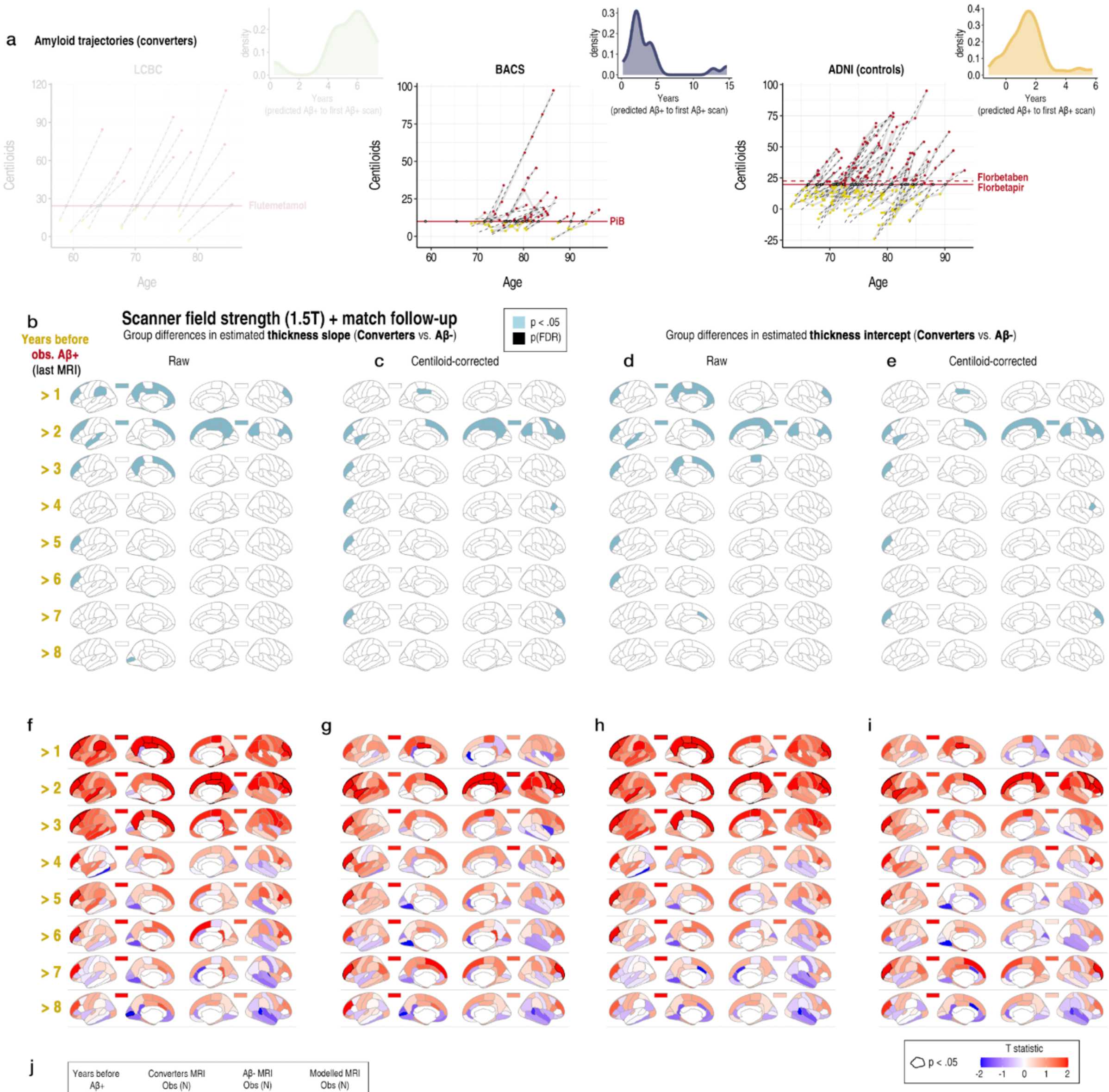

j

| Years before Aβ+ | Converters MRI Obs (N) | Aβ- MRI Obs (N) | Modelled MRI Obs (N) |
|---|---|---|---|
| > 1 | 65 (13) | 107 (26) | 1337 (361) |
| > 2 | 56 (11) | 120 (22) | 1341 (355) |
| > 3 | 50 (10) | 87 (20) | 1302 (352) |
| > 4 | 45 (9) | 106 (18) | 1316 (349) |
| > 5 | 38 (7) | 78 (14) | 1281 (343) |
| > 6 | 31 (6) | 58 (12) | 1254 (340) |
| > 7 | 26 (6) | 48 (12) | 1239 (340) |
| > 8 | 17 (4) | 48 (8) | 1230 (334) |

**Supplementary Figure 18**

**Differences in estimated slope and intercept of cortical thickness between matched converter and Aβ- groups using only 1.5T scans and after discarding the data from LCBC converters (GAMMs). a-c** Amyloid trajectories in centiloids for the Aβ+ converters in each sample (linear model across PET timepoints; raw data in SI Figs. 1-2; see also SI Fig. 13). See Fig. 4. Note that as LCBC only had two PET timepoints, more individuals were predicted to cross the Aβ+ threshold several years before they were observed Aβ+, illustrating how time cutoffs based on years to the first Aβ+ scan should be interpreted with caution. We therefore reran the analysis discarding the data from LCBC converters in the GAMMs. **b-c** Significant differences in the model-estimated slope, with and without adjusting for amyloid levels. **d-e** Significant differences in the model-estimated intercept, with and without adjusting for amyloid.

Black parcels depict FDR-significant hits, whereas blue depicts significance at p<.05 (uncorrected). The rectangles above each brain hemisphere depict the results for mean cortical thickness. **f-i** Corresponding T maps (unthresholded). **i f** The number of MRI observations and N in the mixed-model estimation at each time cutoff, shown for both Aβ groups and the total sample submitted to the GAMMs. Groups were matched on age, and follow-up data.

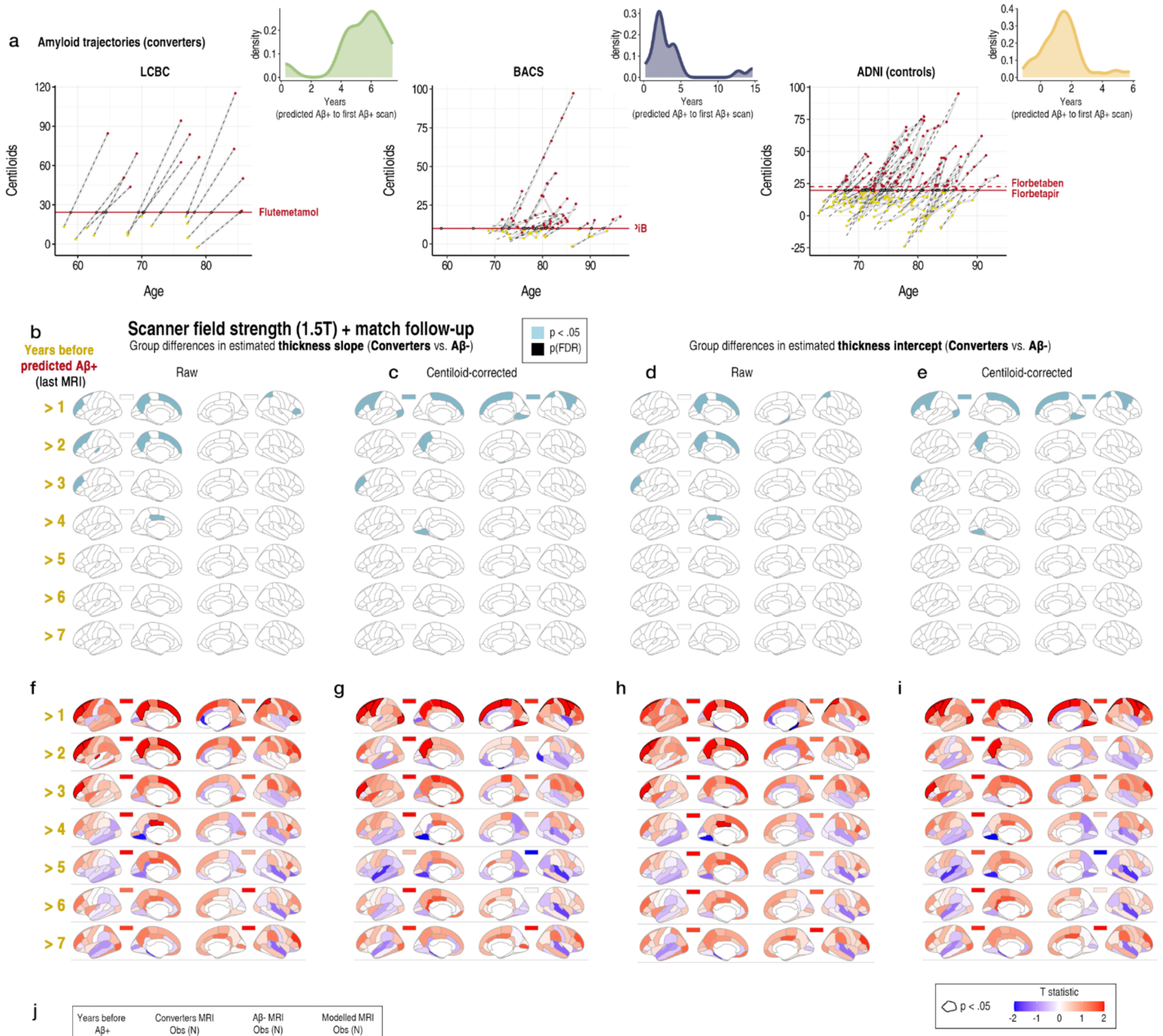

j

| Years before Aβ+ | Converters MRI Obs (N) | Aβ- MRI Obs (N) | Modelled MRI Obs (N) |
|---|---|---|---|
| > 1 | 79 (18) | 160 (36) | 1404 (376) |
| > 2 | 72 (18) | 135 (36) | 1372 (376) |
| > 3 | 61 (16) | 119 (32) | 1345 (370) |
| > 4 | 52 (14) | 107 (28) | 1324 (364) |
| > 5 | 45 (13) | 88 (26) | 1298 (361) |
| > 6 | 36 (11) | 63 (22) | 1264 (355) |
| > 7 | 29 (10) | 55 (20) | 1249 (352) |

**Supplementary Figure 19**

**Differences in estimated slope and intercept of cortical thickness between matched converter and Aβ- groups using predicted age at amyloid positivity (1.5T scans only). a-c** Amyloid trajectories in centiloids (as in main paper). **b-c** Significant differences in the model-estimated slope of cortical thickness, with and without adjusting for amyloid levels (mean centiloids across PET scans prior to the first Aβ$^+$ scan). **d-e** Significant differences in the model-estimated intercept of cortical thickness, with and without adjusting for amyloid. Black parcels depict FDR-significant hits, whereas blue depicts significance at p<.05 (uncorrected). The rectangles above each brain hemisphere depict the results for mean cortical thickness. **f-i** Corresponding T maps (unthresholded). **i f** The number of MRI observations and N in the mixed-model estimation at each time cutoff, shown for both Aβ groups and the total sample submitted to the GAMMs. Groups were matched on age, and follow-up data.

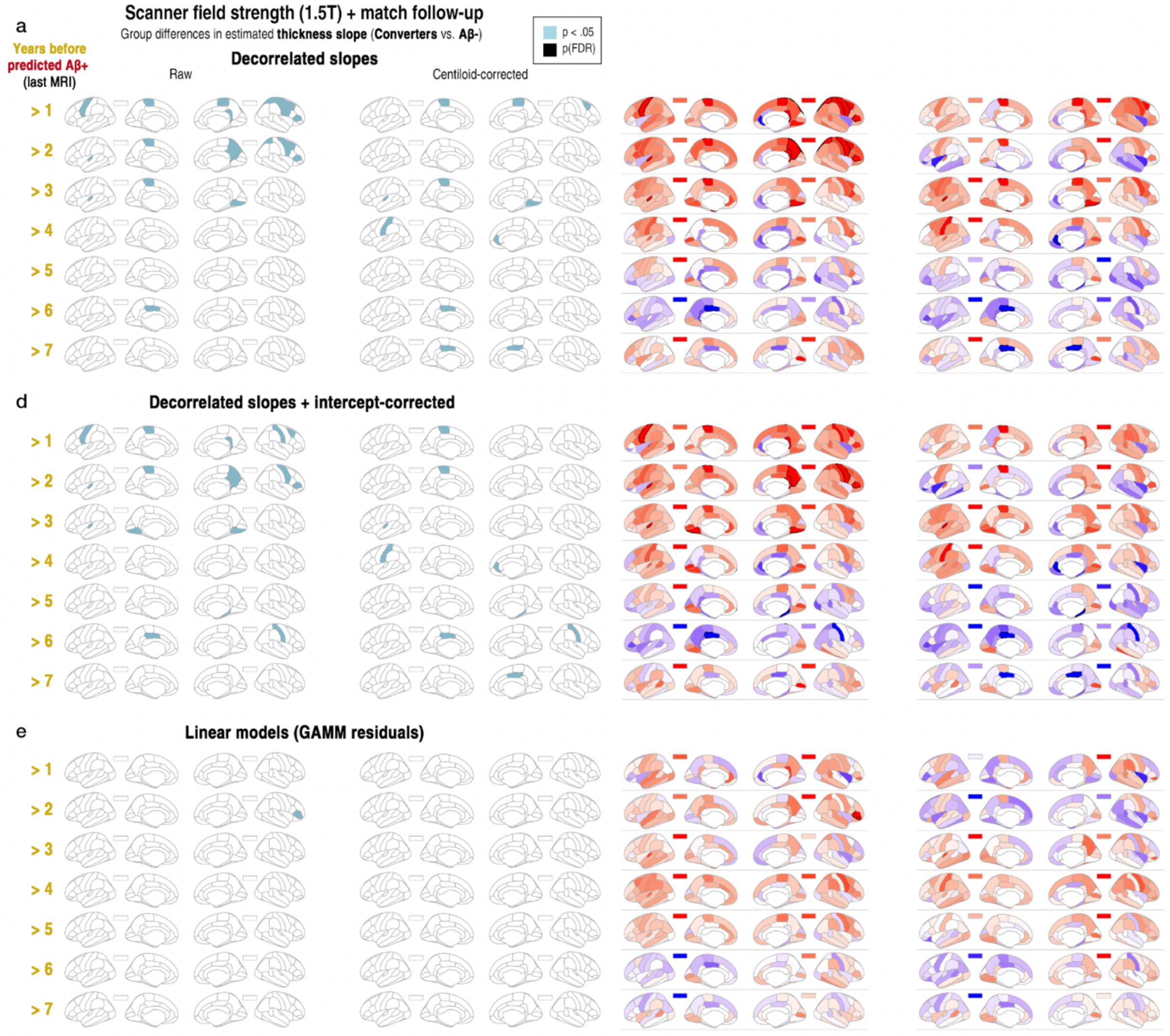

f

| Years before Aβ+ | Converters MRI Obs (N) | Aβ- MRI Obs (N) | Modelled MRI Obs (N) |
|---|---|---|---|
| > 1 | 79 (18) | 160 (36) | 1404 (376) |
| > 2 | 72 (18) | 135 (36) | 1372 (376) |
| > 3 | 61 (16) | 119 (32) | 1345 (370) |
| > 4 | 52 (14) | 107 (28) | 1324 (364) |
| > 5 | 45 (13) | 88 (26) | 1298 (361) |
| > 6 | 36 (11) | 63 (22) | 1264 (355) |
| > 7 | 29 (10) | 55 (20) | 1249 (352) |

**Supplementary Figure 20**
**Differences in cortical thickness change between matched Aβ+ converter and Aβ- groups (models isolating slope from intercept effects using predicted age at amyloid positivity (1.5T scans only). a-b** Significant differences in cortical thickness change between Aβ+ converters and Aβ- groups matched on scanner field strength (1.5T scans only), age and follow-up data. Models shown **a** before and **b** after additionally adjusting for the intercept, and **c** using simple linear models (linear slope across timepoints after regressing out nonlinear age) **f** The number of MRI observations and N in the mixed-model estimation at each time cutoff, shown for both Aβ groups and the total sample submitted to the GAMMs.

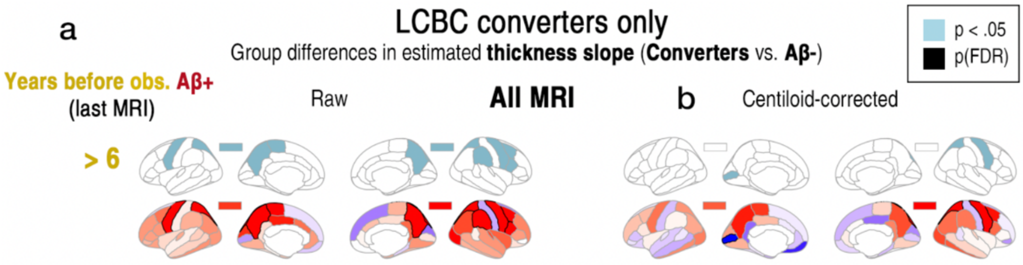

b

| Years before Aβ+ | Converters MRI Obs (N) | Aβ- MRI Obs (N) | Modelled MRI Obs (N) |
|---|---|---|---|
| > 1 | 49 (12) | 95 (24) | 2396 (599) |
| > 2 | 49 (12) | 95 (24) | 2396 (599) |
| > 3 | 49 (12) | 95 (24) | 2396 (599) |
| > 4 | 49 (12) | 95 (24) | 2396 (599) |
| > 5 | 49 (12) | 95 (24) | 2396 (599) |
| > 6 | 49 (12) | 95 (24) | 2396 (599) |

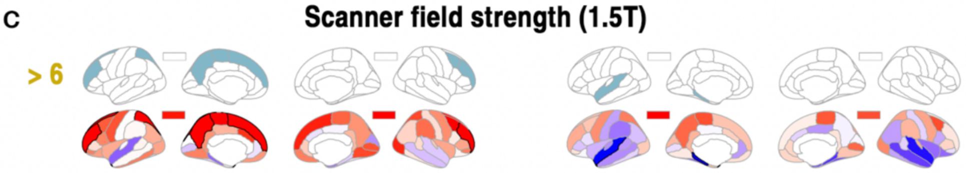

d

| Years before Aβ+ | Converters MRI Obs (N) | Aβ- MRI Obs (N) | Modelled MRI Obs (N) |
|---|---|---|---|
| > 1 | 26 (9) | 53 (18) | 1244 (349) |
| > 2 | 26 (9) | 53 (18) | 1244 (349) |
| > 3 | 26 (9) | 53 (18) | 1244 (349) |
| > 4 | 26 (9) | 53 (18) | 1244 (349) |
| > 5 | 26 (9) | 53 (18) | 1244 (349) |
| > 6 | 26 (9) | 53 (18) | 1244 (349) |

**Supplementary Figure 21**
**Differences in estimated slope of cortical thickness between matched converter and Aβ- groups after discarding the data from BACS and ADNI converters (LCBC converters only).**

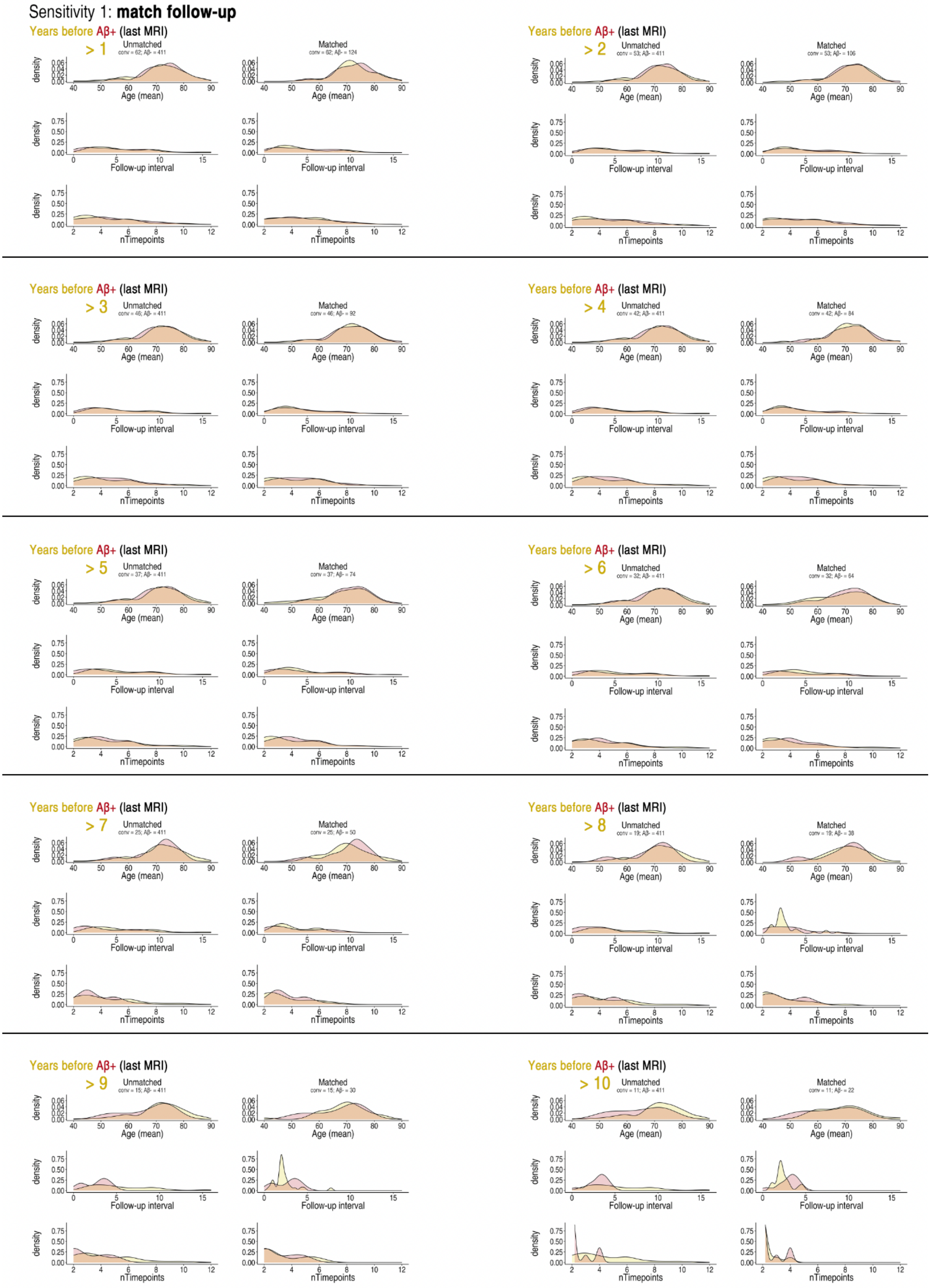

**Supplementary Figure 22**

Matching groups on follow-up data. At each time cut-off we drew a matched sample from the Aβ- group, based on mean age, the number of MRI timepoints, and the time interval covered in the converter group. Because the N of the converter group was smaller (max N = 62 [1 year cutoff]), we repeated the matching twice, drawing two one-to-one matched samples from the Aβ- group.

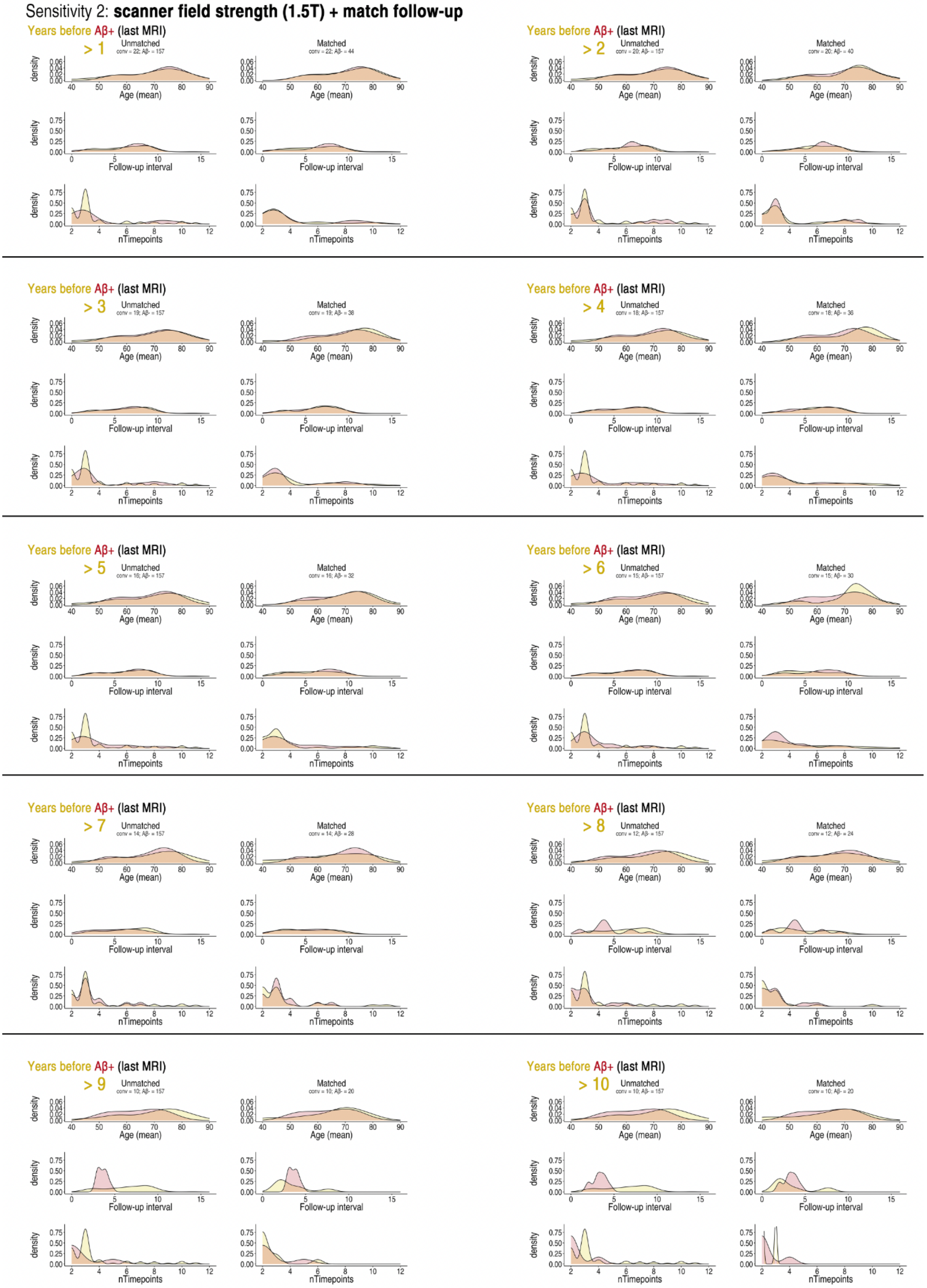

**Supplementary Figure 23**
Matching groups on scanner field strength (1.5T) and follow-up data. At each time cut-off we drew a matched sample from the Aβ- group, based on mean age, the number of MRI timepoints, and the time interval covered in the converter group. Because the N of the converter group was smaller (max N = 22 [1 year cutoff + 1.5T scans only]), we repeated the matching twice, drawing two one-to-one matched samples from the Aβ- group.

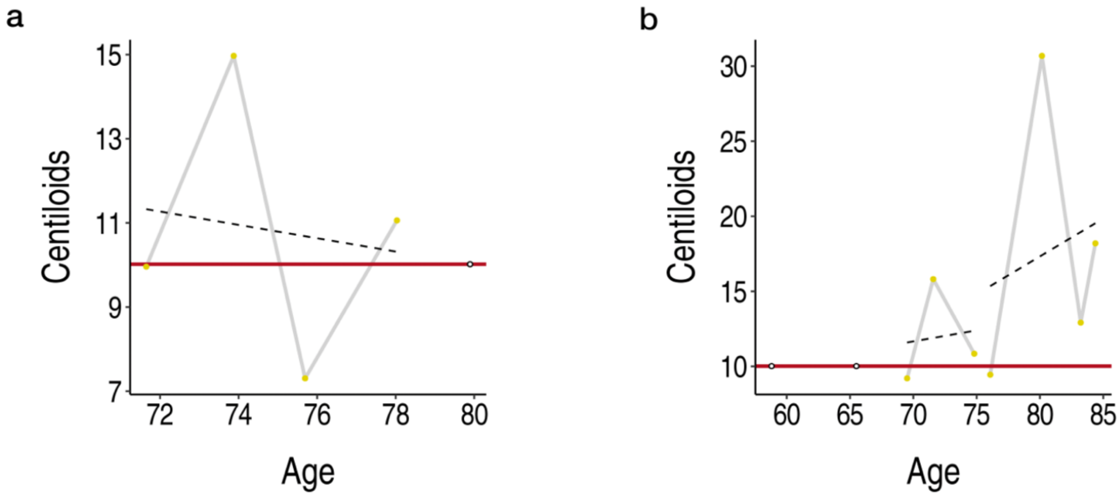

**Supplementary Figure 24**
**a** For analyses using predicted age at amyloid positivity**,** one converter was excluded as their centiloid slope over time was negative (despite their unambiguous Aβ+ status), such that their predicted age on crossing the sample-specific Aβ+ threshold in centiloids was too high (white dot). **b** Two converters were predicted to cross the threshold at very low ages, likely attributable to measurement error in their centiloid trajectories.

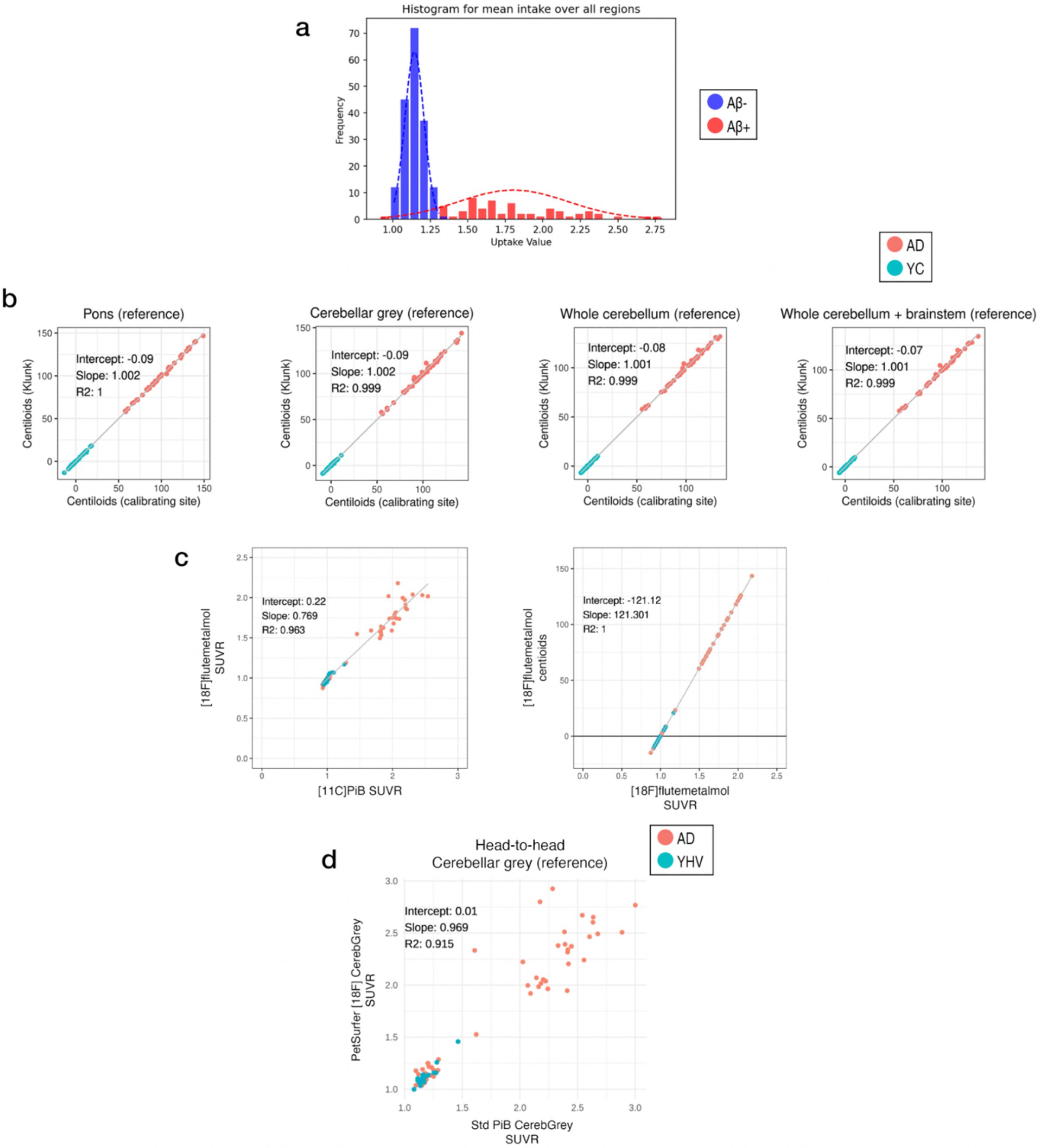

**Supplementary Figure 25**

**Aβ modelling and centiloid calibration with GAAIN pipeline (LCBC [18F]flutametamol-PET scans). a** Results of gaussian mixed modelling to define Aβ status in LCBC data. Note the one individual with SUVR<1.0 estimated on the Aβ+ distribution was set to Aβ-. **b** Replication of published centiloid values using standard pipeline. The calculated anchorpoint group means were 1.014 ($YC_0$) and 2.086 ($AD_{100}$) (whole cerebellum reference). Results were within the expected individual (±5%) and group (±2%) tolerances and expected agreement with published results ($r^2$ >.99) in Klunk et al.[3] **c** Replication of correlation between 18F SUVR units (standard pipeline) and [11C]PiB SUVR units (standard pipeline) in head-to-head AD-control GAAIN data (‘GE_flutemetamol.xlsx’; https://www.gaain.org/centiloid-project). **d** Correlation between PiB SUVR units (standard pipeine) and 18F SUVR units using site-specific pipeline (PetSurfer) in head-to-head AD-control GAAIN data. Estimation of PiB SUVR units from 18F SUVR units derived from PetSurfer pipeline: PiB = ($SUVR_{18F}$ – 0.006) / 0.969. Centiloid equation: CL = 100*(PiB – $YC_0$) / ($AD_{100}$ – $YC_0$).

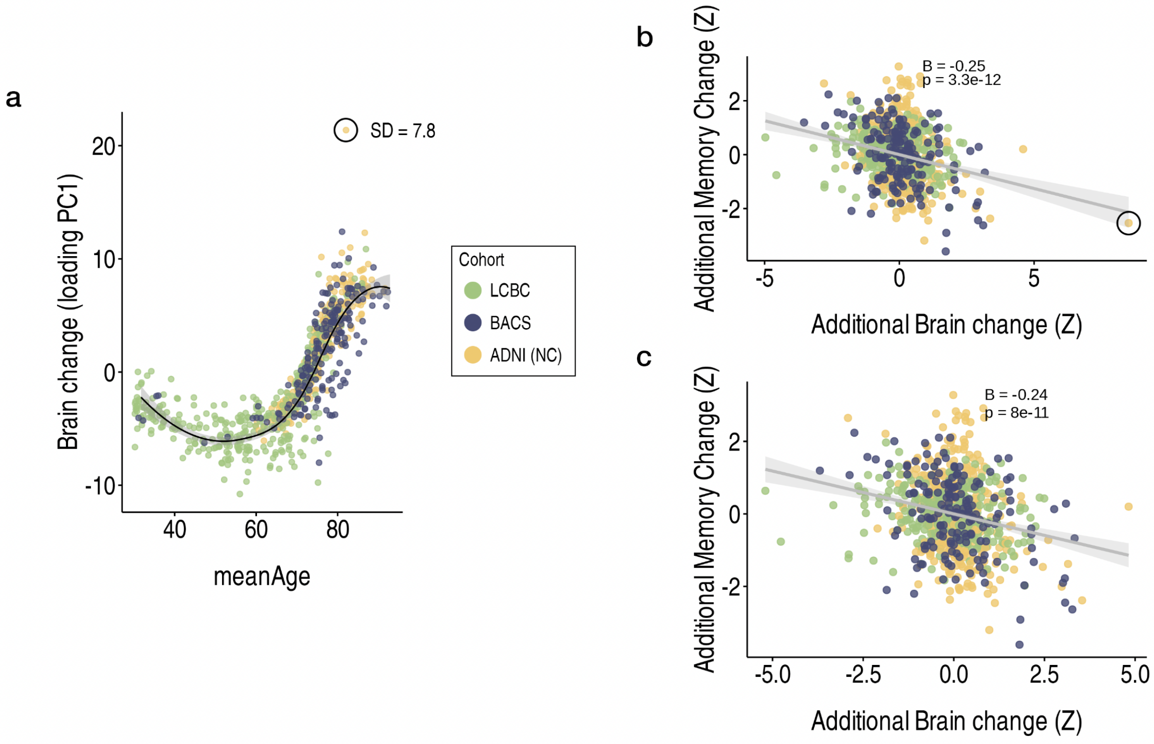

**Supplementary Figure 26**
**ADNI outlier. a** One strong outlier (SD = 7.8) in the brain change data of ADNI controls was identified and removed from all MRI analyses. **a** The outlier was detected via the principal component (PC1) of absolute change across the top 50 brain features with accelerated change in AD (see Fig. 4 in [7]). We removed this outlier from all MRI analyses due to unrealistic change values. **b-c** Note that their inclusion strengthened (rather than attenuated) the association between brain and memory decline in cognitively healthy adults (unpublished results). The outlier was an Aβ- subject, but we did not test how their inclusion affected the results of the present paper.

**Supplementary Note 1**

For LCBC, all subjects underwent a [$^{18}$F]Flutemetamol amyloid-PET scan to assess cortical Aβ burden[8]. Images were acquired on a GE HealthCare Discovery PET/CT 690 scanner at Aleris Hospital and Radiology, Oslo, Norway. PET images were processed using PetSurfer, the MRI-PET analysis tool within the FreeSurfer (v7.1.0) package. The intensities were processed with Region-Based Voxel-wise Partial Volume Correction (RBV-PVC) and scaled by the cerebellar cortex (left and right). The resulting uptake values were then further analyzed with the sklearn (v1.5.1) toolbox. Principal component analysis was used to extract the first principal component (PC1) from the standardized uptake value ratios (SUVR) in the region of interest described by Mormino et al.[9]. Next, assuming data normality, PC1 was used to estimate the parameters of the Gaussian Mixture Model (GMM) with two mixture components; thus, two distributions were estimated. This allowed the probability of each observation belonging to each distribution to be calculated. Those falling under the distribution with the higher mean were classified as A+ participants.

**Supplementary Note 2**

**Dataset access**

Requests for access to the raw data can be directed to the Principal Investigators of the contributing studies: LCBC (Anders M. Fjell;). BACS, data are available on request to William J. Jagust. ADNI (https://adni.loni.usc.edu/data-samples/access-data/).
